# Volumetric Two-photon Imaging of Neurons Using Stereoscopy (vTwINS)

**DOI:** 10.1101/073742

**Authors:** Alexander Song, Adam S. Charles, Sue Ann Koay, Jeff L. Gauthier, Stephan Y. Thiberge, Jonathan W. Pillow, David W. Tank

**Affiliations:** Department of Physics, Princeton University; Princeton Neuroscience Institute, Princeton University; Bezos Center for Neural Circuit Dynamics, Princeton University; Department of Psychology, Princeton University; Department of Molecular Biology, Princeton University

## Abstract

Two-photon laser scanning microscopy of calcium dynamics using fluorescent indicators is a widely used imaging method for large scale recording of neural activity in vivo. Here we introduce volumetric Two-photon Imaging of Neurons using Stereoscopy (vTwINS), a volumetric calcium imaging method that employs an elongated, V-shaped point spread function to image a 3D brain volume. Single neurons project to spatially displaced image pairs in the resulting 2D image, and the separation distance between images is proportional to depth in the volume. To demix the fluorescence time series of individual neurons, we introduce a novel orthogonal matching pursuit algorithm that also infers source locations within the 3D volume. We illustrate vTwINS by imaging neural population activity in mouse primary visual cortex and hippocampus. Our results demonstrate that vTwINS provides an effective method for volumetric two-photon calcium imaging that increases the number of neurons recorded while maintaining a high frame-rate.

## Introduction

Two-photon excitation laser scanning microscopy (TPM) [1] enables high spatial resolution optical imaging in highly scattering tissue such as the mammalian brain. When combined with genetically-encoded calcium indicators [2, 3], or synthetic indicators that label neural populations [4], intracellular calcium dynamics can be measured across a population of cells, providing a method for large scale recording of neural activity at cellular resolution [4, 5]. In general, increasing the number of simultaneously recorded neurons is important because it increases the power of population analysis methods in studies of neural coding and dynamics. To increase the number of neurons recorded with two-photon calcium imaging, volumetric imaging methods, such as multi-plane imaging [6], random access fluorescence microscopy [7-9] and ultrasound lens scanning [10], are under development.

In traditional TPM [1], a Gaussian excitation beam, focused to a diffraction-limited spot, is scanned in a raster pattern across the sample and an image is created from measurement of the emitted fluorescence at each location. The non-linear process of fluorophore excitation, together with the sharp axial falloff in intensity in the point spread function (PSF), leads to optical sectioning: the image from one raster scan represents fluorescence intensity in one plane within a sample volume. To increase the number of recorded neurons beyond those resolved in a single plane, volume imaging can be performed by sequentially moving the focal plane (or sample) up or down between each raster scan, repeating this pattern for each volume measurement. This method can be implemented with movable objectives, remote focusing [11], or a liquid lens [6]. However, if the frame rate for single plane imaging is *N* frames/sec, and the number of planes imaged per volume in *m*, then the aggregate volume frame rate is reduced to *N*/*m*. Many calcium indicators have on-response kinetics below 0.1 s [12]. To capture this dynamics, volume frame rates must remain close to 10 Hz. With current resonant scanner-based TPM (*N* ≈ 30 Hz), this implies that only a relatively low number of planes (m=3,4) can be used for multi-plane volumetric imaging.

Elongating the PSF of the focused excitation beam along the optical axis, using either a low-NA Gaussian beam focus or Bessel beam methods [13], can be used with raster scanning to form a projection image of a volume [14]. This is useful in applications like functional imaging of dendritic spines in sample volumes with sparse neural expression of the indicator [15]. However, in samples with dense expression, such as those encountered in large-scale recording of a neural population in vivo, extending a single PSF axially causes neurons at different depths to be superimposed. Information about depth in the sample of individual neurons is lost, and demixing of fluorescence signals from individual neurons is compromised if their images signi cantly overlap.

Our method addresses these limitations by using an elongated PSF that is split into two excitation beams. These beams are spatially separated and angled inwards to create a stereoscopic “V”-shaped PSF configuration (Fig. 1a). Raster scanning with this PSF produces a 2D projection image that preserves information about neural activity at different depths. We refer to this method as *volumetric Two-photon Imaging of Neurons using Stereoscopy* (vTwINS). The intuition behind vTwINS is straightforward: the soma of any neuron in the 3D volume makes two contributions to the 2D projection image, one soma-shaped image for each arm of the V-shaped PSF. The spatial offset between these two images is equal to the distance between the two arms of the V at the neuron's depth in the volume. This results in short distances between deep neurons, and longer distances for shallower neurons (Fig. 1a).

**Figure 1.**
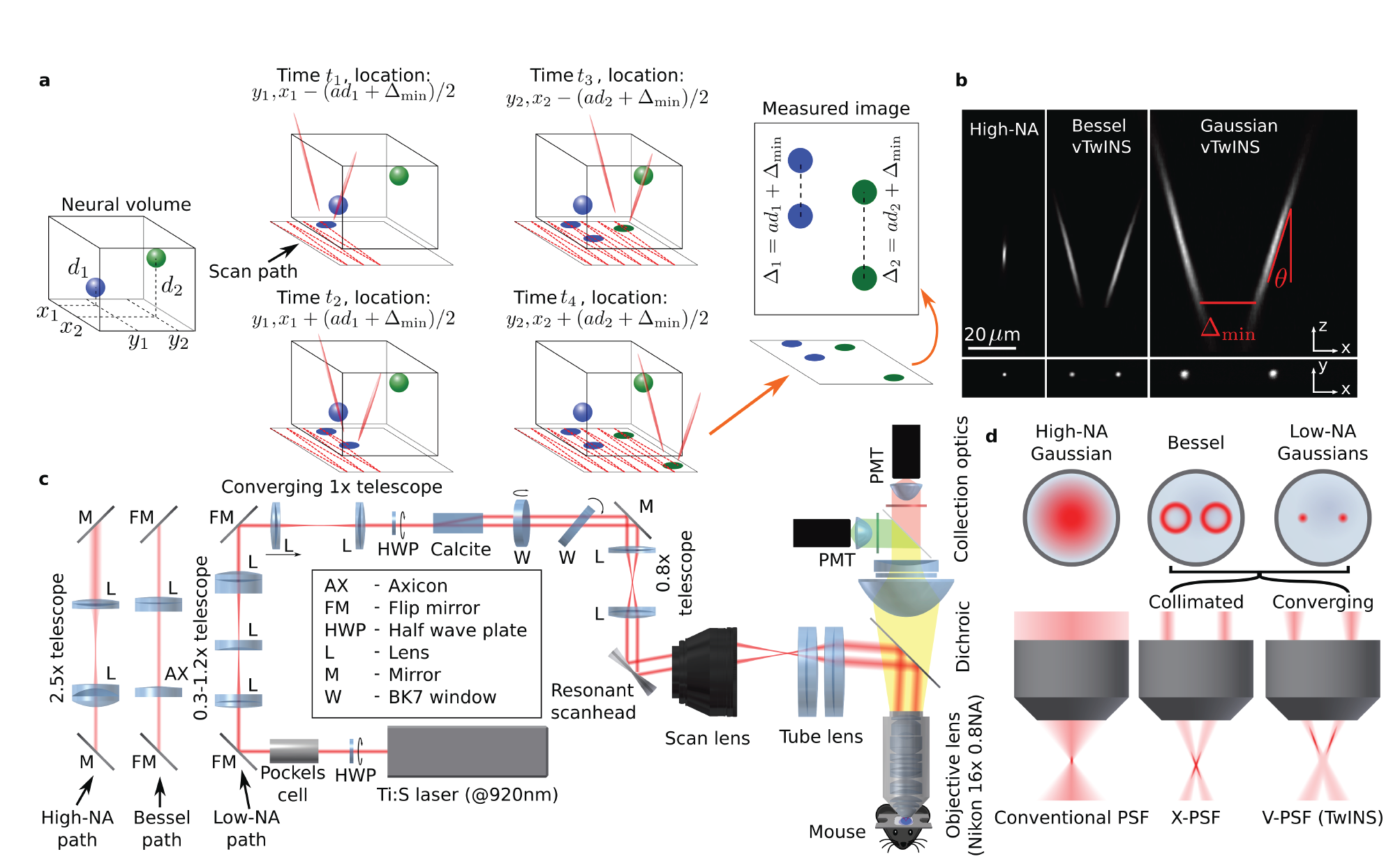
vTwINS concept and design. (a) vTwINS uses a “V”-shaped PSF to image neural volumes. During scanning, the two PSF arms intersect neurons at different depths (e.g. the blue and green stylized neurons) with different time intervals. Deep neurons intersect the second arm shortly after the first. Shallow neurons take longer for the second arm to intersect. Each neuron thus appears twice, where the distance between images indicates depth. (b) Example PSFs for diffraction-limited (high-NA) TPM, and vTwINS microscopes using Bessel and low-NA Gaussian beams. (c) The vTwINS microscope consists of a beam-shaping module and a conventional two-photon microscope. The three optical paths generate the PSFs shown in (b). In the Bessel and Gaussian (low-NA vTwINS paths, lenses adjust the PSF's axial extent, and a birefringent block (calcite) splits the beam in two and sets the PSF angle. (d) The back aperture illumination profiles for the three paths in (c). In the high-NA (conventional TPM) path, the overfilled back aperture is focused to a point. In the Bessel and low-NA Gaussian paths, two beams are focused to form each arm of the PSF. The beam divergence is adjusted with the 1x telescope before the calcite block to separate the two arms of the X-PSF and form the V-PSF.

Although vTwINS ensures that all neurons will have distinct paired spatial profiles in the projection image, the analysis problem of identifying which image regions correspond to pairs reflecting the activity of single neurons is ill-posed from single images. This diiffculty can be solved by a demixing algorithm that relies on the temporal statistics of neural activity across frames. There are many approaches to spatial profile identification and demixing from time series, including ICA/PCA [16] and constrained non-negative matrix factorization (CNMF) [17, 18]. Our approach was to extend prior work to the case where the expected shape of the neuron's spatial profile is a pair of rings or disks displaced along the axis of the V-shaped PSF. We describe a novel inference algorithm based on orthogonal matching pursuit that exploits both the spatial separation of image pairs and the sparseness of neural activity. The algorithm's identification of a neuron's spatial profile in the projection image also means that a neuron's relative depth *d* in the volume scanned can be recovered from the image pair separation Δ via the relationship *d*=0.5(△–△_min_/tan(θ)), where △_min_ is the minimum inter-beam distance of the PSF and θ is the beam angle from the axial direction (Fig. 1a,b). Thus, the demixing algorithm both provides the time course of fluorescence change and information from which the neuron's location in the volume can be reconstructed.

In the following, we describe the optics developed to produce the vTwINS PSF and demonstrate images and image time series produced using this method. We then present the algorithm that was developed for identifying active neurons in these time series and demixing fluorescence transients. Finally, using the combined imaging system and algorithm, we demonstrate large-scale recording of GCaMP-expressing neurons in visual cortex and hippocampus of the awake mouse.

## Results

### vTwINS Optics

In a vTwINS microscope the diffraction limited PSF (Fig. 1b left) of a traditional raster scanning TPM is replaced with an elongated V-shaped PSF produced from two intersecting Gaussian beams (Fig. 1b center), or Bessel beams (Fig. 1b right). The strategy we used to create the V-shaped PSF was dual beam excitation through a single objective lens (Fig. 1c), using the design principles illustrated in Figure 1d. In the traditional TPM, a large diameter collimated Gaussian beam is centered on the objective back aperture, and a standard (typically diffraction-limited) PSF for two-photon excitation is produced (Fig. 1d left). In contrast, a pair of smaller diameter collimated Gaussian beams with their centers offset from the center of the back aperture produce a pair of elongated arms in the PSF that cross at the focal plane, producing an X-shape (Fig. 1d center). The smaller the diameter of each beam, which reduces the effective NA, the more each arm becomes elongated. Increasing the separation between the two beams at the objective back aperture increases the angle of intersection of the two arms. If the incident beams are slightly divergent (or convergent), the position of the crossing point of the two beams shifts along the optical axis relative to the focal point of the objective (Fig. 1d right), eventually producing a V-shape for vTwINS, with the wider opening either pointing up (divergent beams) or down (convergent beams). To produce a Bessel beam vTwINS PSF, rings of illumination are used at the objective back aperture [19] instead of Gaussian beams, but, otherwise, the same principles apply. Bessel beams allow for more uniform axial excitation, at the cost of excitation eiffciency (see Discussion).

To explore vTwINS imaging with either Bessel or Gaussian profiles, and to compare the images to those generated by a standard PSF (diffraction-limited single-beam), we designed a beam-shaping module (Fig. 1c) with three parallel beam paths, one for each modality, with a set of flip mirrors that could select between them. A calcite block was used to split a single incident beam into the two spatially separated beams necessary for vTwINS. Input polarization, controlled by a half-wave plate, was used to equalize power in each of the two beams in the vTwINS configuration, and to eliminate beam splitting when a standard single-beam PSF was used. The angle between the two arms of the PSF in vTwINS mode was determined by the beam separation produced by the calcite block together with a subsequent telescope. Beam divergence, used to control where the two beams cross (and forming either a V or inverted V), was controlled by an adjustable telescope placed before the beam splitter. The arm used for Bessel beam vTwINS differed from that of the Gaussian beams by replacing the first telescope with an axicon-lens combination that formed the ring-shaped spatial profile required for Bessel beam production.

As an initial proof of principle that a set of fluorescence sources could be spatially localized in a 3D volume from a single vTwINS image, we imaged a volume sample of 1 *μ*m diameter fluorescent latex beads embedded in agar. The beads were embedded at random locations, creating an off-grid set of positions. The exact bead positions were determined via a diffraction-limited two-photon multi-plane volumetric scan (z-stack) using the traditional PSF. vTwINS was then used to image the same volume with a single scan (one image) using a 58 *μ*m-long PSF (FWHM, 75 *μ*m 1/e full-width). Each bead produces a pair of dots in the vTwINS projection image (Supplementary Fig. 1); lines drawn between all pairs are parallel and aligned with the direction of the vTwINS PSF in the sample. The distance between dot pairs varies with the bead's depth in the volume. Using only the vTwINS image and the known shape of the PSF to infer each bead's 3D coordinates in the sample produces average errors of 1.4±1.3 *μ*m in depth, 1.5±1.3 *μ*m in the fast-scan direction, 1.2±1.0 *μ*m in the slow-scan direction and an average total localization error of 2.7±1.6 *μ*m. The accuracy of recovered positions is well within the ≈10 *μ*m average size of a neuronal cell body in the mammalian brain, demonstrating that vTwINS, in practice, preserves the necessary information to disambiguate cell bodies at different depths.

**Supplementary Figure 1:**
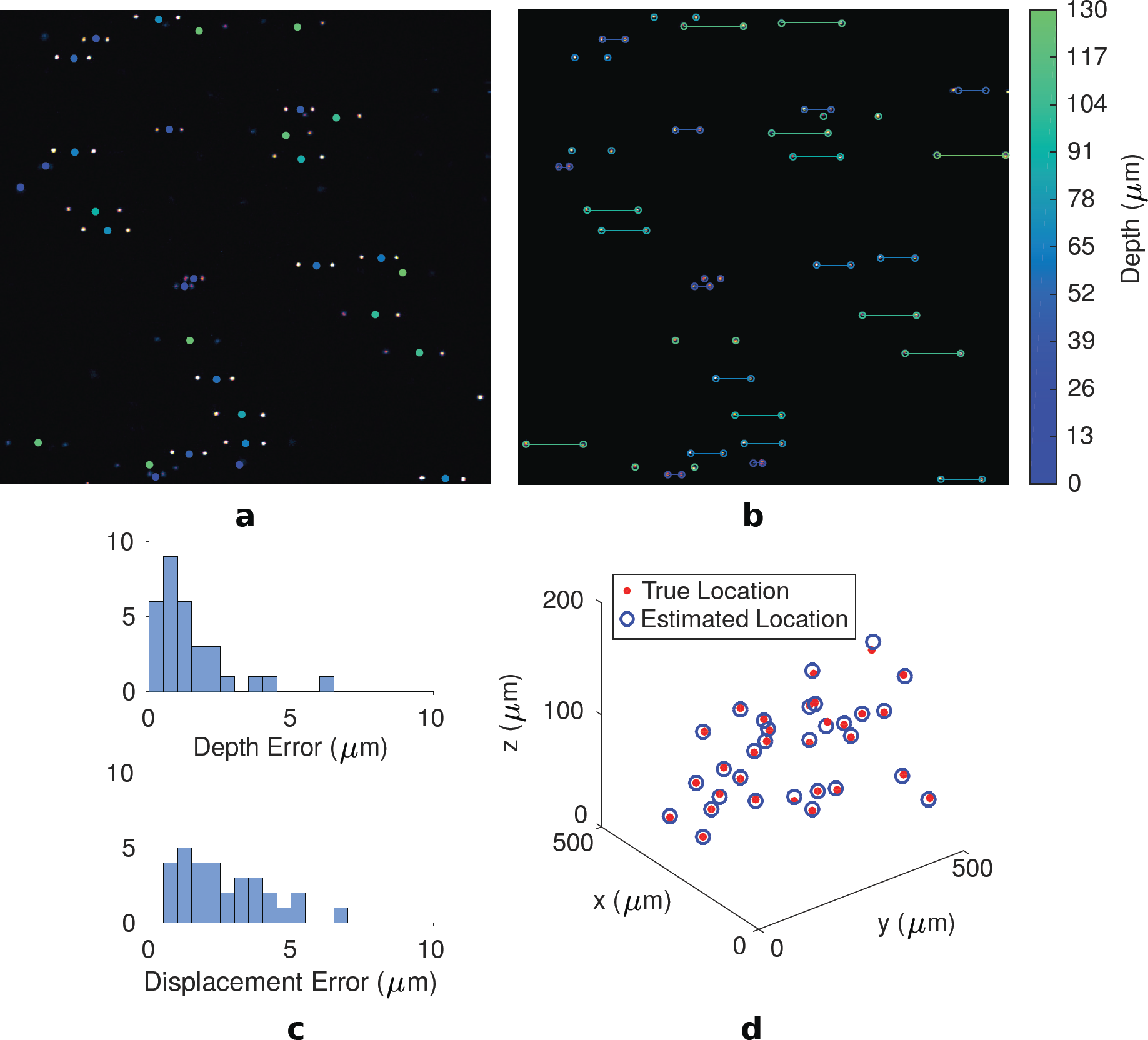
vTwINS depth recovery of fluorescent beads. (a) vTwINS image of a volume containing beads at different locations. Circles depicting locations of all the beads in the vTwINS axial range are color coded by depth (blue is deeper, indicating nearer images, and green is shallower, implying wider images). (b) Spatial profiles recovered automatically from the single vTwINS image are color coded on the same depth scale. Beads at the edge of the field of view have occluded images, and are thus excluded from the analysis as depth cannot be ascertained. (c) Histograms depicting the axial localization error in as well as the total displacement error show that vTwINS can recover bead locations to within approximately 5µm. (d) The 3D scatter plot compares both the true location of the beads (red dots) and the estimated location (blue circles) to better visualize the vTwINS accuracy.

### vTwINS Calcium Imaging

The basic features of vTwINS-based calcium imaging data, obtained from visual cortex (V1) in an awake transgenic mouse expressing GCaMP6f (see Methods), are illustrated in Figure 2.. A single image plane taken with TPM using a diffraction-limited PSF is also shown for comparison. In diffraction-limited TMP (Fig. 2a), a single soma-shaped spatial profile of high fluorescence intensity is observed when calcium transients are produced in an active neuron. The cell soma of some, but typically not all [20], GCaMP-expressing quiescent cells can also be resolved. A vTwINS image is qualitatively different. Active neurons in a vTwINS image become represented as two bright soma shaped regions (disk or ring; Fig. 2b). Additionally, the images of quiescent neurons are typically not resolved because the projection produces an increased, and more uniform, background intensity, which is due to the axially extended PSFs exciting a larger volume of tissue that includes neuropil and other cell bodies. When multiple cells are simultaneously active, many soma pairs become visible, and lines drawn between corresponding image pairs are parallel and oriented along the fast scan direction (the plane of the V-shaped PSF). However, pairs from different cells can have different spatial separations (Fig. 2c), representing different depths of the cell somas in the volume. A time series vTwINS movie from mouse V1 is provided in Supplementary Video 1.

**Figure 2:**
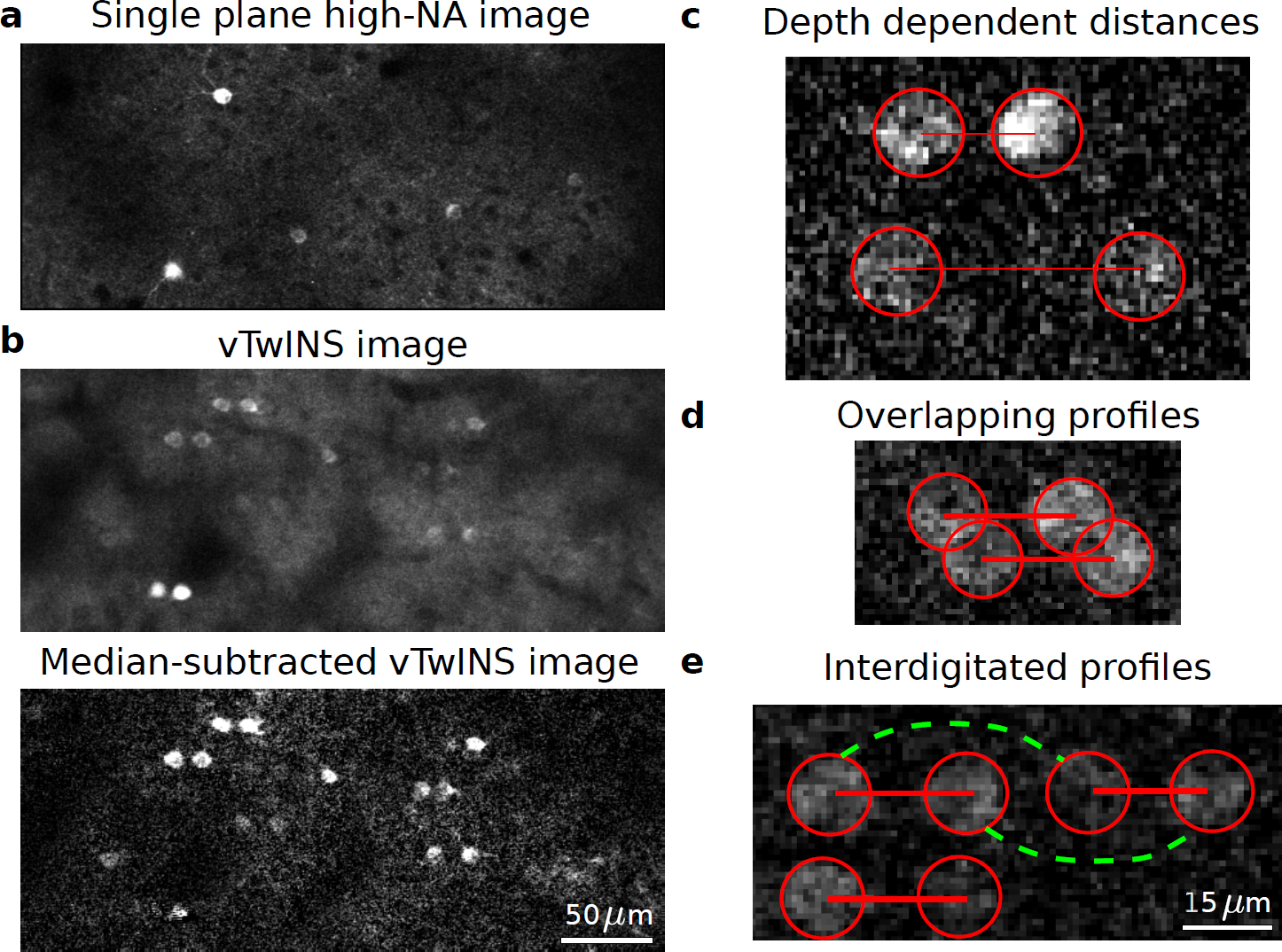
Example vTwINS images. All images are averages of 5 consecutive frames. (a) Diffraction-limited TPM single plane image of GCaMP in mouse visual cortex. (b) vTwINS scan of the same V1 area as (a) demonstrates paired somas of active neurons and reduced SNR as the background levels are much higher. Subtracting the temporal median at each pixel highlights neural activity. (c) Two fluorescing neurons imaged by vTwINS at different depths have different distances between the image pairs. Red circles indicate the different images and red lines connect corresponding image pairs. (d) vTwINS images typically have overlapping spatial profiles. (e) Neurons aligned in the direction parallel to the plane of the V PSF (which is the same as the fast scan direction in our implementation) can create ambiguity in the spatial profile image pair assignment. Both the solid red lines (the true pairing) and the dashed green lines indicate realizable distance pairings corresponding to different neuron positions, and temporal activity must be used to resolve this ambiguity.

The properties of vTwINS based calcium imaging data (Fig. 2) introduce a number of unique challenges in demixing spatial profiles of neural activity in order to extract the time traces of fluorescence for individual cells. First, there is a lower SNR per cell due to the axially extended PSFs (Fig. 2b). Second, the spatial profiles of cells under vTwINS can partially overlap (Fig. 2d), and typically consist of disjoint regions, violating the spatial locality assumption in current demixing methods [16,18]. Third, neurons co-aligned in the fast-scan direction can create ambiguous, interdigitated spatial profile pairs (Fig. 2e). Finally, intensity differences between the two images in a pair may result from the non-uniform scattering between the two beam paths (e.g. due to varying tissue properties).

### vTwINS profile identification and Demixing

We addressed the challenges of analyzing vTwINS data with Sparse Convolutional Iterative Shape Matching (SCISM), a novel demixing method that explicitly seeks horizontally separated image pairs. A graphical summary of the method is provided in Figure 3, while the specifics and the full statistical model are provided in the Methods. As a pre-processing step we motion-corrected, temporally averaged and spatially binned the raw image time series (see Methods). At each iteration, candidate spatial profiles, consisting of stereotyped profiles (designed as pairs of annuli separated in the fast-scan direction with different inter-image distances; Fig. 3a), are compared to the measured fluorescence frames across the field-of-view (FOV) (Fig. 3b). The stereotyped profile most correlated with the data is then selected (Fig. 3c), and the most highly correlated frames are used (Fig. 3d) to refine the profile shape to better match the data. This step allows SCISM to handle spatial profile pairs where one beam path has lower intensity. The new profile is added to the set of spatial profiles, and the corresponding time-traces are estimated via a non-negative LASSO [21] procedure (Fig. 3e). Finally, the data residual is calculated by subtracting the component of the data captured by the current set of spatial profiles (Fig. 3f), and this residual is reused in the correlation step to determine the next spatial profile.

**Figure 3:**
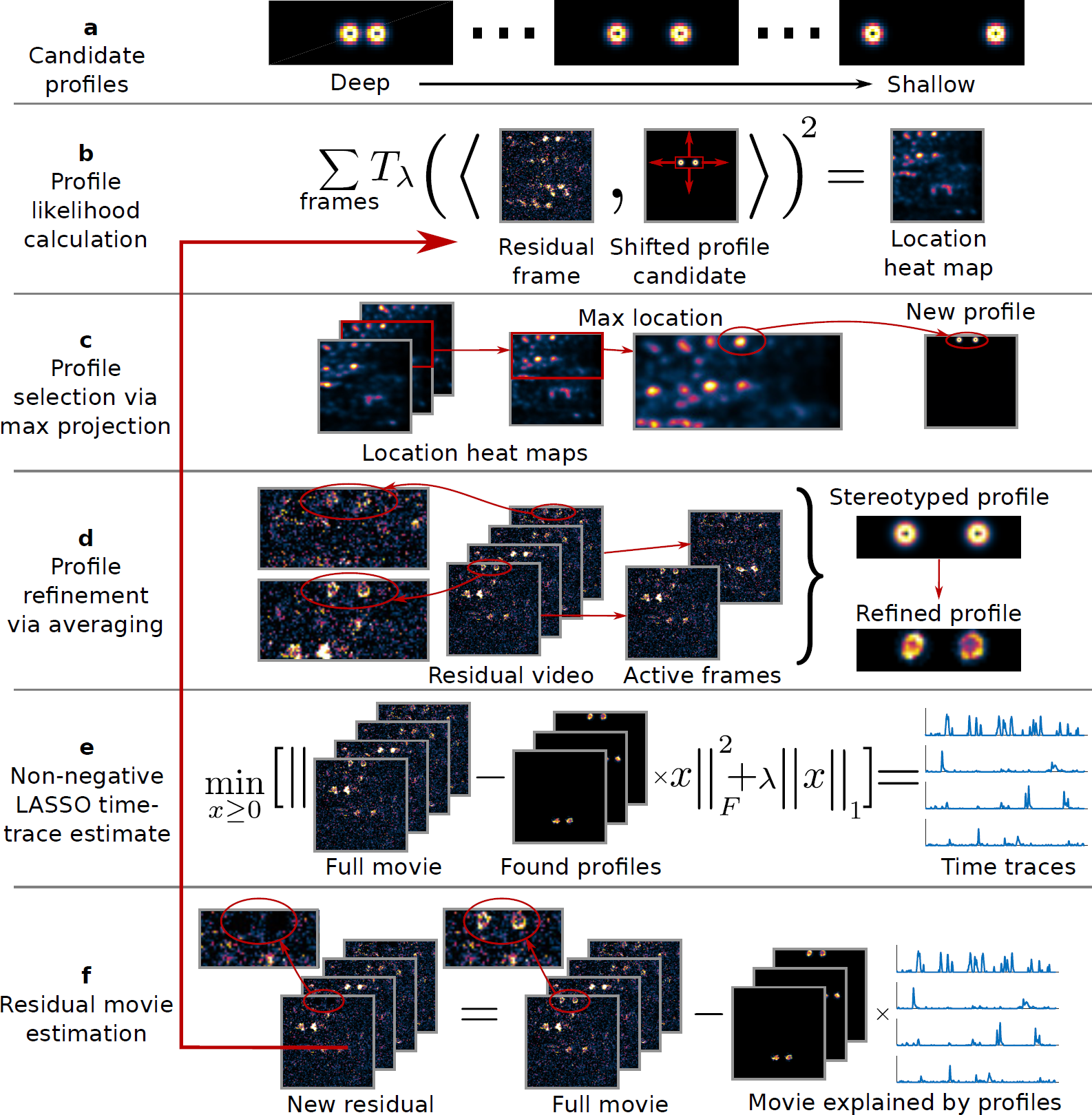
Sparse convolutional iterative shape matching (SCISM) for demixing vTwINS data. (a) Example stereotyped neuron image pairs. (b) SCISM seeks image pairs at different distances by constructing heat-maps representing the likelihood of a given pair at a given location. Heat maps are calculated by summing the thresholded squared-inner-product between shifts of stereotyped profiles and video frames (shown here with a section of CA1 data). *T_λ_*(.) here denotes the threshold operation. (c) The new spatial profile is chosen at the maximum across all heat maps. (d) The new profile is refined by locally masking and averaging frames closely aligned with the stereotyped spatial profile. (e) The new profile is added to the set of spatial profiles, and the time-traces for all spatial profiles are calculated via non-negative LASSO with sparsity trade-off parameter λ (see Methods). (f) The residual movie is re-computed by subtracting the contribution of the current set of spatial profiles (the sum of outer products of the spatial profiles and their time traces). The algorithm then finds the next spatial profile by iterating from (b) with the new residual.

This procedure iteratively selects spatial profiles greedily in order of correlation strength with the data, using both spatial and temporal statistics to determine the most likely spatial profile at each iteration. Specifically, SCISM leverages sparsity in neural activity as well as the spatial constraint that each spatial profile consists of two areas separated in the fast-scanning direction. Sparse neural activity is particularly important as it permits minimal cross-contamination due to spatially overlapping neurons. Once spatial profiles are determined with SCISM, full resolution time trace estimates are obtained using non-temporally averaged data via non-negative LASSO.

In the following, we demonstrate the application of SCISM to vTwINS calcium imaging data from both visual cortex (V1) and hippocampus (CA1) in awake transgenic mice expressing GCaMP6f. For V1 recordings we performed full FOV vTwINS imaging and we validated our results against diffraction-limited TPM by performing simultaneous imaging with the two modalities for half-size FOV. For CA1 we demonstrated the utility of vTwINS in neural volumes with a very high density of neurons.

### Large Scale Recording in Mouse Visual Cortex

Head-restrained GCaMP6f-expressing transgenic mice, running on a spherical treadmill, were presented with a visual stimulus sequence consisting of randomly placed Gabor patches (see Methods).

vTwINS imaging was performed in layer 2/3 of primary visual cortex (V1). Images were acquired in a 550 *μ*m × 550 *μ*m area with a 45 *μ*m-long inverted-V PSF (FWHM, 60 *μ*m 1/e full-width) at 30 Hz frame rate over a 14 minute imaging session.

The time series fluorescence data was preprocessed with rigid motion-correction and spatio-temporal averaging (see Methods, Supplementary Fig. 2, and Supplementary Video 1,2). Spatial profiles obtained via SCISM (Fig. 4) show significant overlap, as expected from the relatively high density of GCaMP-expressing cells and the vTwINS PSF. Given the spatial profiles, we used the vTwINS PSF to extract the 3D cell positions (see Methods, Fig. 4a,b). The demixed spatial profile activity traces (Fig. 4c, Supplementary Fig. 3, Supplementary Fig. 4) have the expected temporal statistics of sparsely firing neurons. Because SCISM is an iterative method that extracted highly active spatial profiles first, the time traces are ordered by how correlated the profiles are with the data.

**Figure 4:**
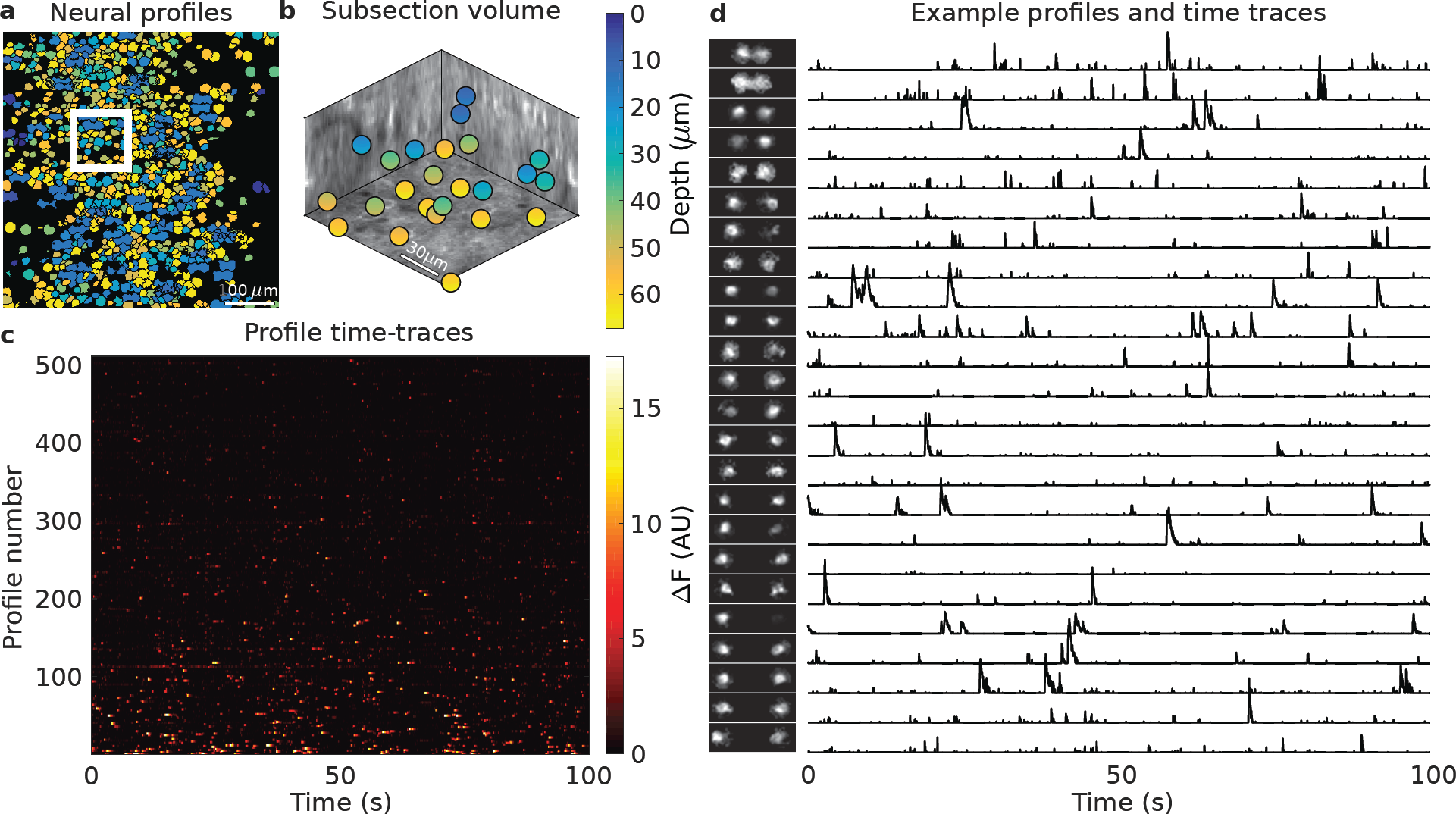
Demixed spatial profiles and calcium activity in mouse visual cortex. (a) Full set of spatial profiles, color-coded by depth, show significant overlap. (b) 3D locations of the spatial profiles located in the white box in (a) show that spatial profiles are found at different depths. (c) Time-traces of spatial profiles show sporadic activity in the 0-100s time interval. (d) Example subset of spatial profiles (chosen from the white inset box in (a) and sorted by depth) and corresponding time traces show rich activity patterns. The increasing separation distance as a function of depth reflects the inverted V shape of the PSF used in this recording.

**Supplementary Figure 2:**
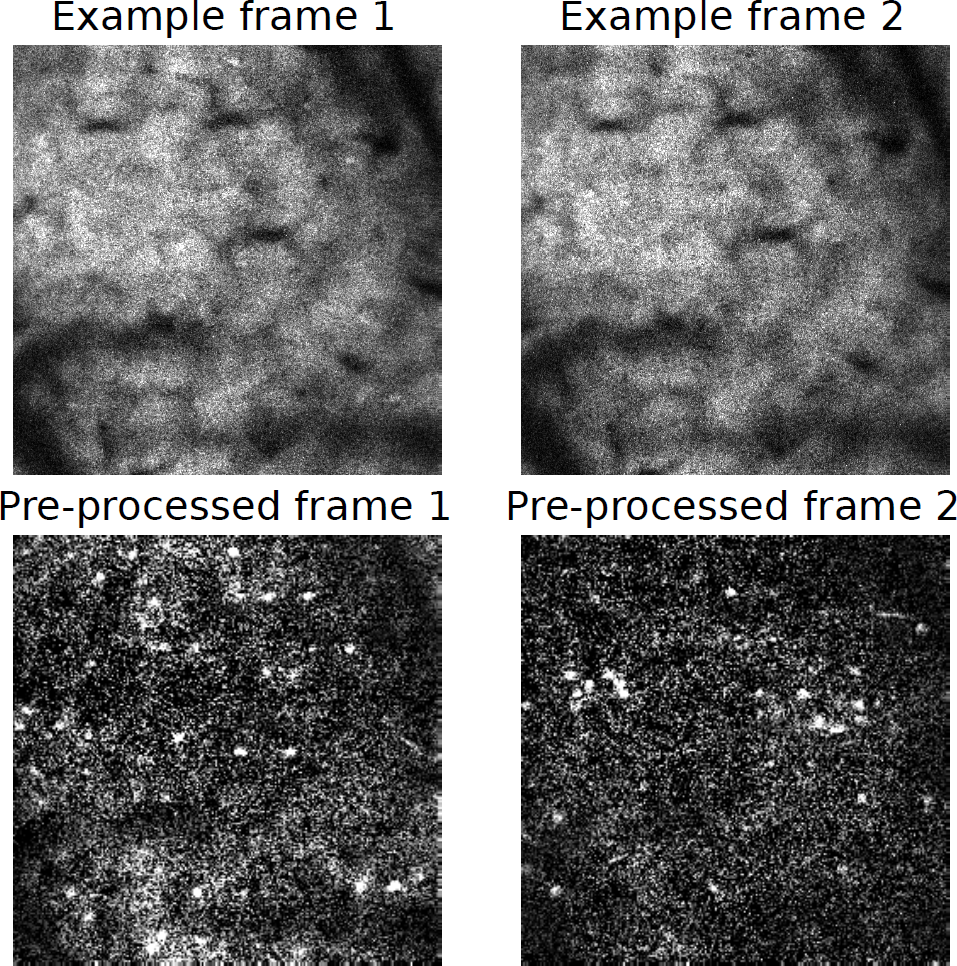
Example frames of full FOV vTwINS data acquired from V1. Top: two examples of vTwINS images from V1. Bottom: Corresponding pre-processed images (5-frame temporal average and two-fold spatial binning) with background-subtraction. Pre-processing makes active pairs of neuronal images more apparent.

**Supplementary Figure 3:**
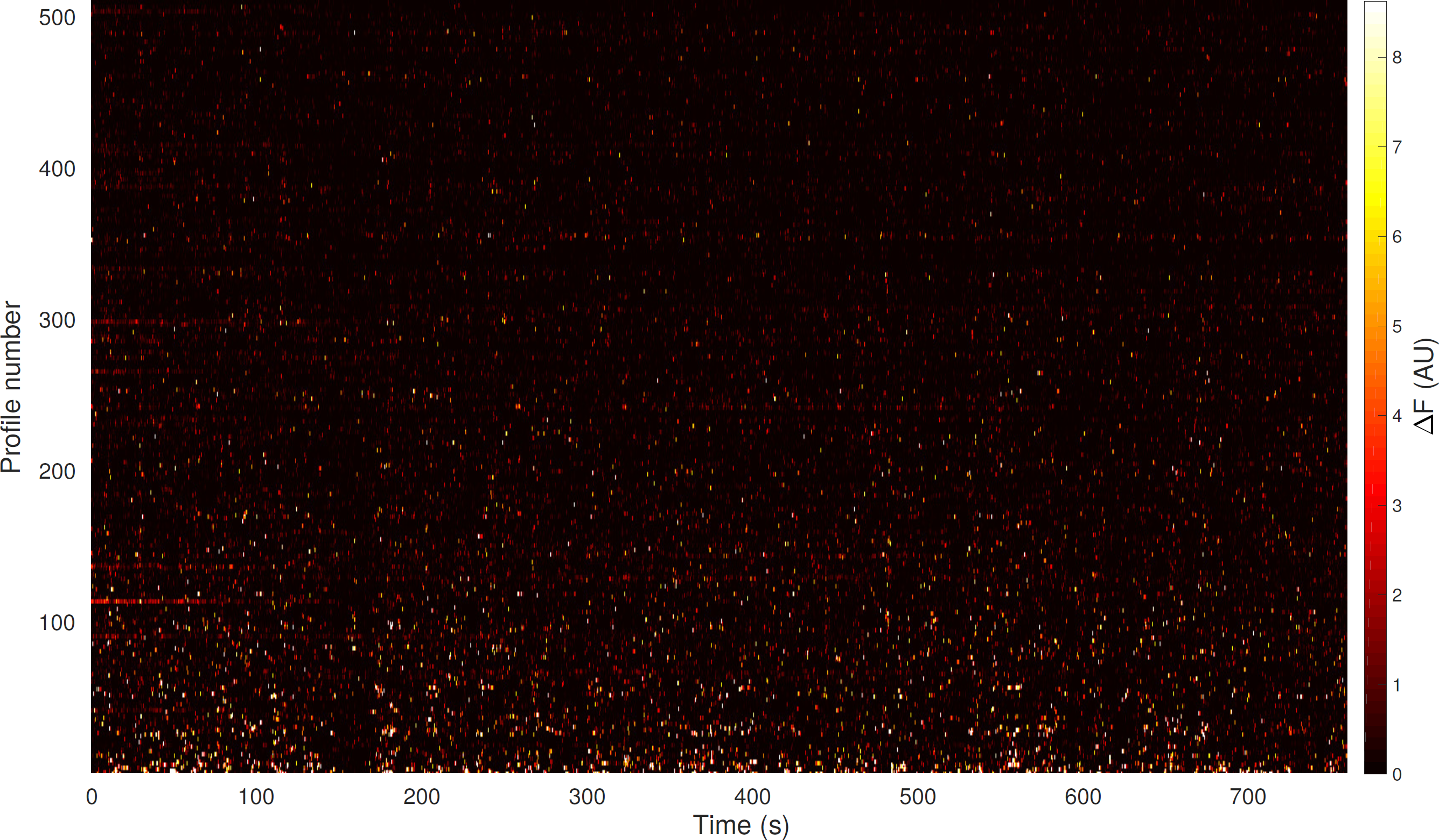
800 second time traces for all demixed V1 spatial profiles.

**Supplementary Figure 4:**
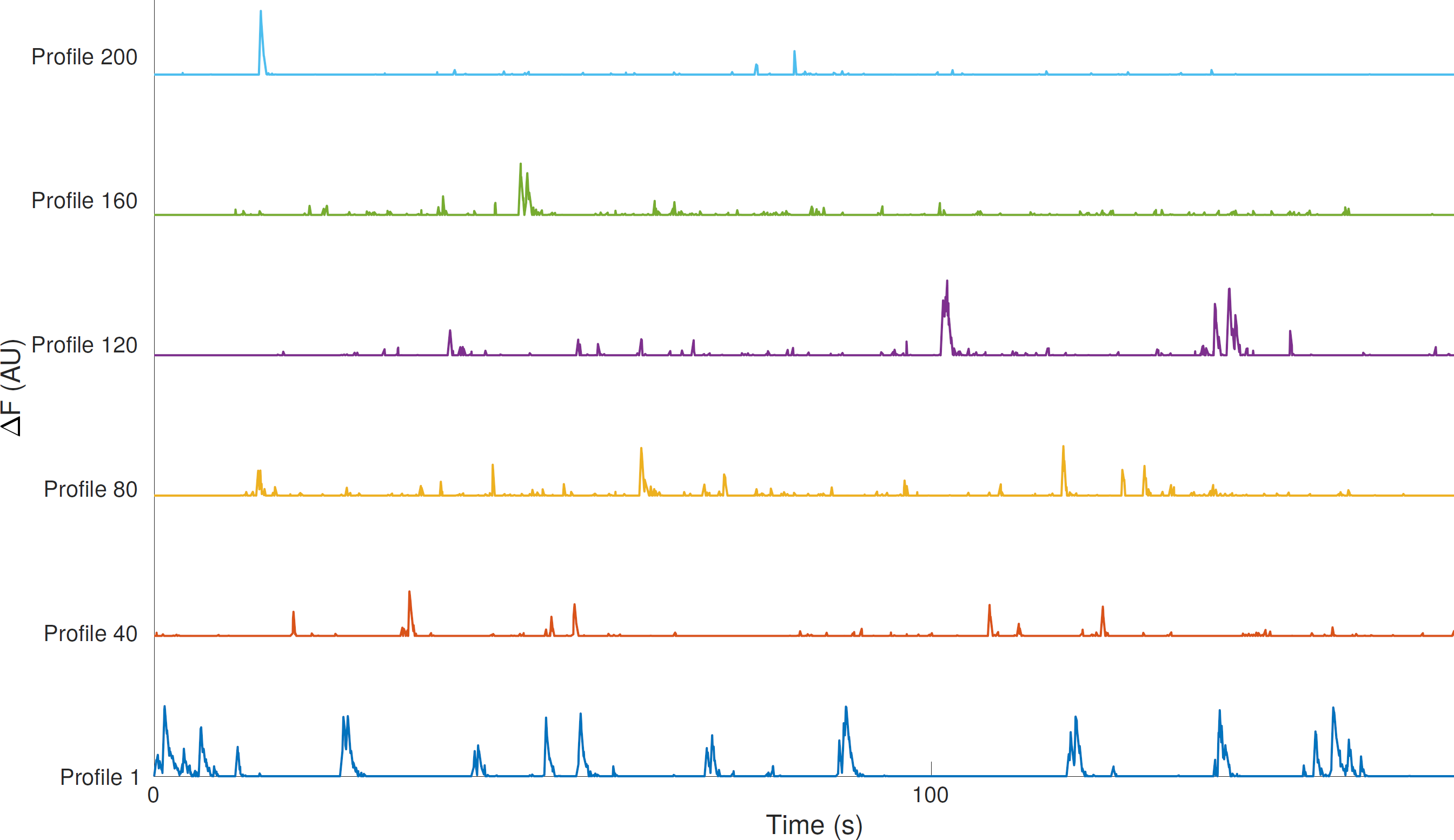
Example time traces for select demixed V1 spatial profiles.

The spatial profile volumetric locations (Fig. 4b) indicates that vTwINS records activity across the entire axial extent. The range of axial depths captured by vTwINS is further illustrated by plotting the spatial profiles in a 107 *μ*m × 107 *μ*m subsection of the FOV (Fig. 4d), sorted by inferred depth, (Fig. 4d) and their corresponding position in a 3D anatomical volume (see Methods, Supplementary Fig. 5). We note that all the spatial profiles in this subsection have a clear cell body in the anatomical z-stack, which corresponds to the calculated 3D position. The large variation in separation distance between the spatial profile image pairs suggests that vTwINS does capture and demix activity originating from many axial planes. The corresponding spatial profile activity traces (Fig. 4d) also show that cell transients are well isolated, despite the highly overlapping spatial profiles.

**Supplementary Figure 5:**
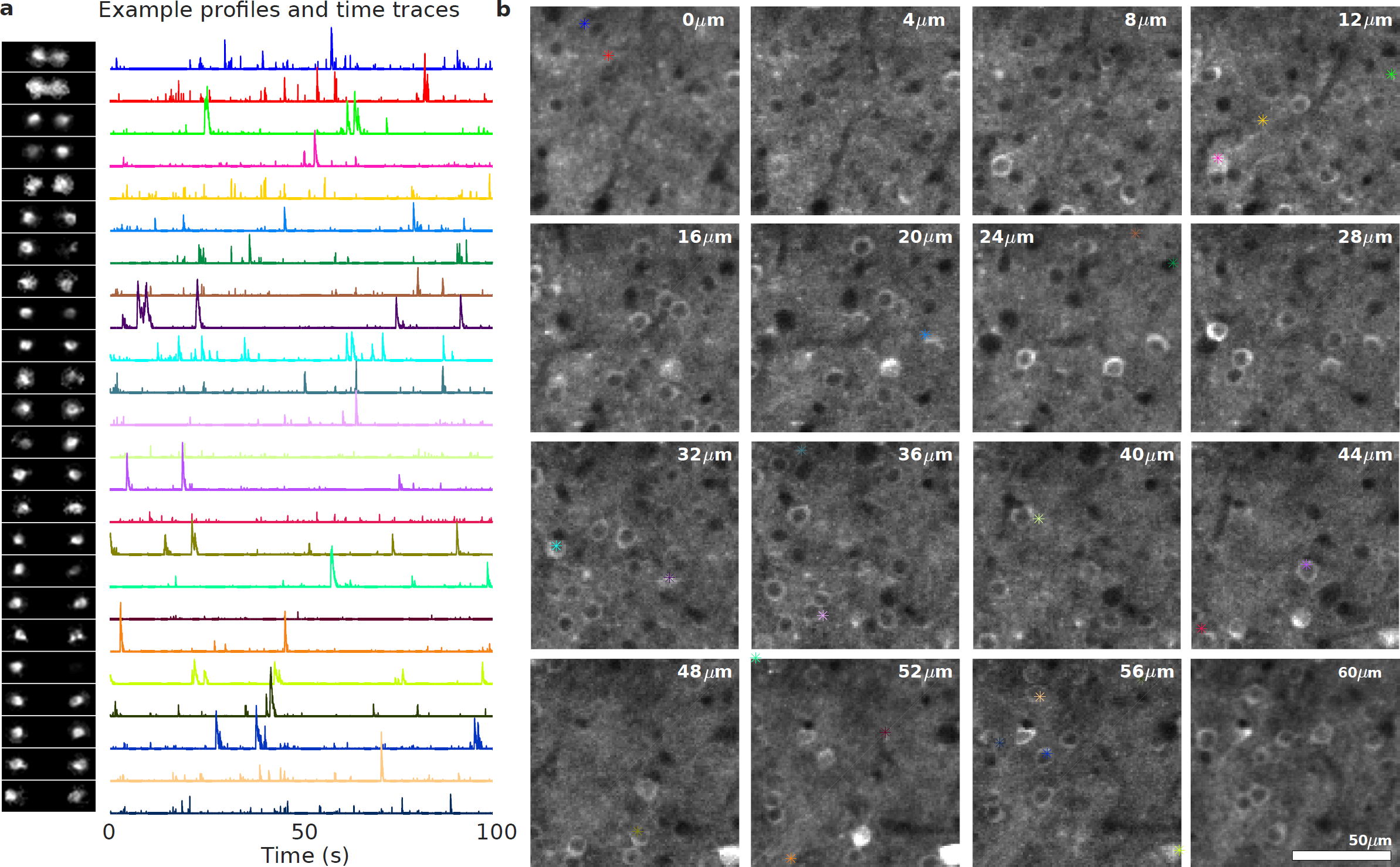
Anatomical z-stack comparison of found spatial profiles in V1. (a) Example spatial profiles and time traces from V1. (b) Corresponding location of activity in the anatomical z-stack. White numbers at the top of each image indicates the relative depth in the z-stack.

To validate that neural activity recorded with vTwINS is comparable to a standard method, we compared vTwINS with single-plane diffraction-limited TPM. We verified both spatial profile locations and temporal activity by simultaneously imaging an entire neural volume with vTwINS, and a single slice of the volume with diffraction-limited TPM. Both datasets were collected at 30 Hz over a 470 *μ*m × 200 *μ*m overlapping area. A galvanometer was used to flip between the vTwINS, using a 38 *μ*m-long PSF (FWHM, 52 *μ*m 1/e full-width), and standard TPM beampath configurations at every frame (see Supplementary Fig. 6). Alternating between imaging modalities at every scan resulted in two interleaved movies recording the same neural activity with a ≈17 ms offset between corresponding frames. Aside from introducing the interleaving mechanism, the only other difference in this recording from the full FOV V1 recording (Fig. 4) was that the vTwINS PSF used was the noninverted “V”-shaped PSF. vTwINS data was demixed using SCISM (see Methods, Fig. 3), while we extracted spatial profiles and activity traces from the single-plane data using a modified constrained non-negative matrix factorization (CNMF) algorithm [22] as an independent comparison (see Methods). Spatial profiles from the two methods were identified as arising from the same neuron using both the centroid locations and the level of temporal correlation in the demixed activity traces (see Methods).

**Supplementary Figure 6:**
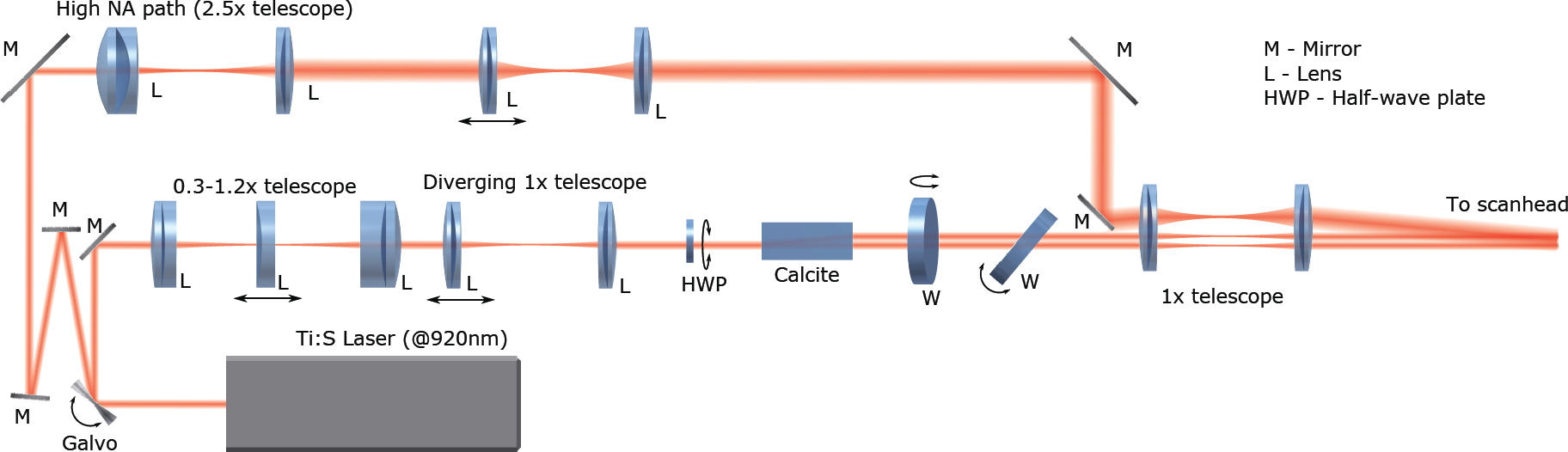
Alternate optical setup for simultaneous imaging using vTwINS and conventional TPM. After each frame, a galvanometer switches between a conventional high-NA TPM path and a low-NA Gaussian vTwINS path. A mirror is used to recombine the two paths, with an offset angle of 0.88°.

Comparison of spatial profiles from the simultaneous recordings (Fig. 5) indicates that vTwINS captures both neural activity in the single slice TPM and activity at other depths. To highlight examples of correlated cells, a 5 s max-projection was used to select a fraction of active cells within the volume (Fig.5a,b). Overall, in a ten minute imaging session, 454 spatial profiles were found in the volume using vTwINS, as compared to 169 spatial profiles found in the single plane diffraction-limited data. Activity traces corresponding to the found spatial profiles of cells identified in both the single plane and the volume also show very high correlation between the two imaging modalities (Fig. 5c). This correlation indicates that vTwINS still captures most of the activity at any given depth while also capturing the additional activity elsewhere in the volume. Specifically, of the single-slice spatial profiles, 116 spatial profiles had >1 transient per minute. Of these, 98 (84%) had a matching spatial profiles in the vTwINS data (Supplementary Fig. 7). Of the remaining single-slice spatial profiles, many had very low SNR, suggesting that that activity fell below the vTwINS' lower SNR level.

**Figure 5:**
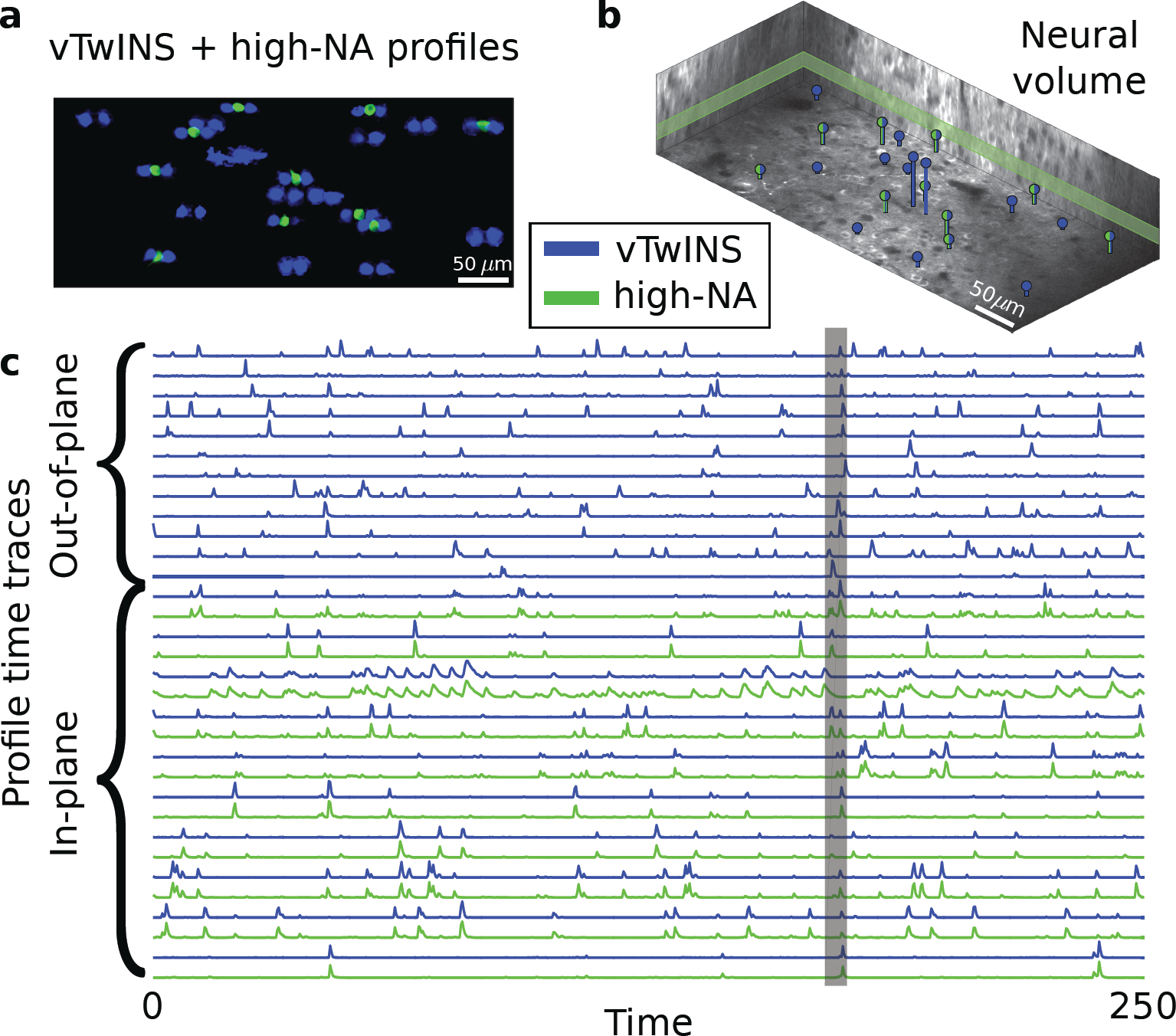
Simultaneous imaging of visual cortex using conventional two-photon (green) and vTwINS (blue). (a) Max-projection of 5 s of activity (top) and corresponding extracted spatial profiles (bottom) demonstrate that the spatial profile extraction algorithms demix relevant neural activity. (b) Volumetric depiction of vTwINS extracted spatial profile locations and depth. Green/blue profiles indicate location of cells that were matched in single-plane activity. The single-plane slice is outlined in green. (c) Time traces corresponding to cells in (a) show that for vTwINS with diffraction-limited TPM counterparts, the temporal activity traces match. The gray bar indicates the 5s period of activity used to isolate cells.

**Supplementary Figure 7:**
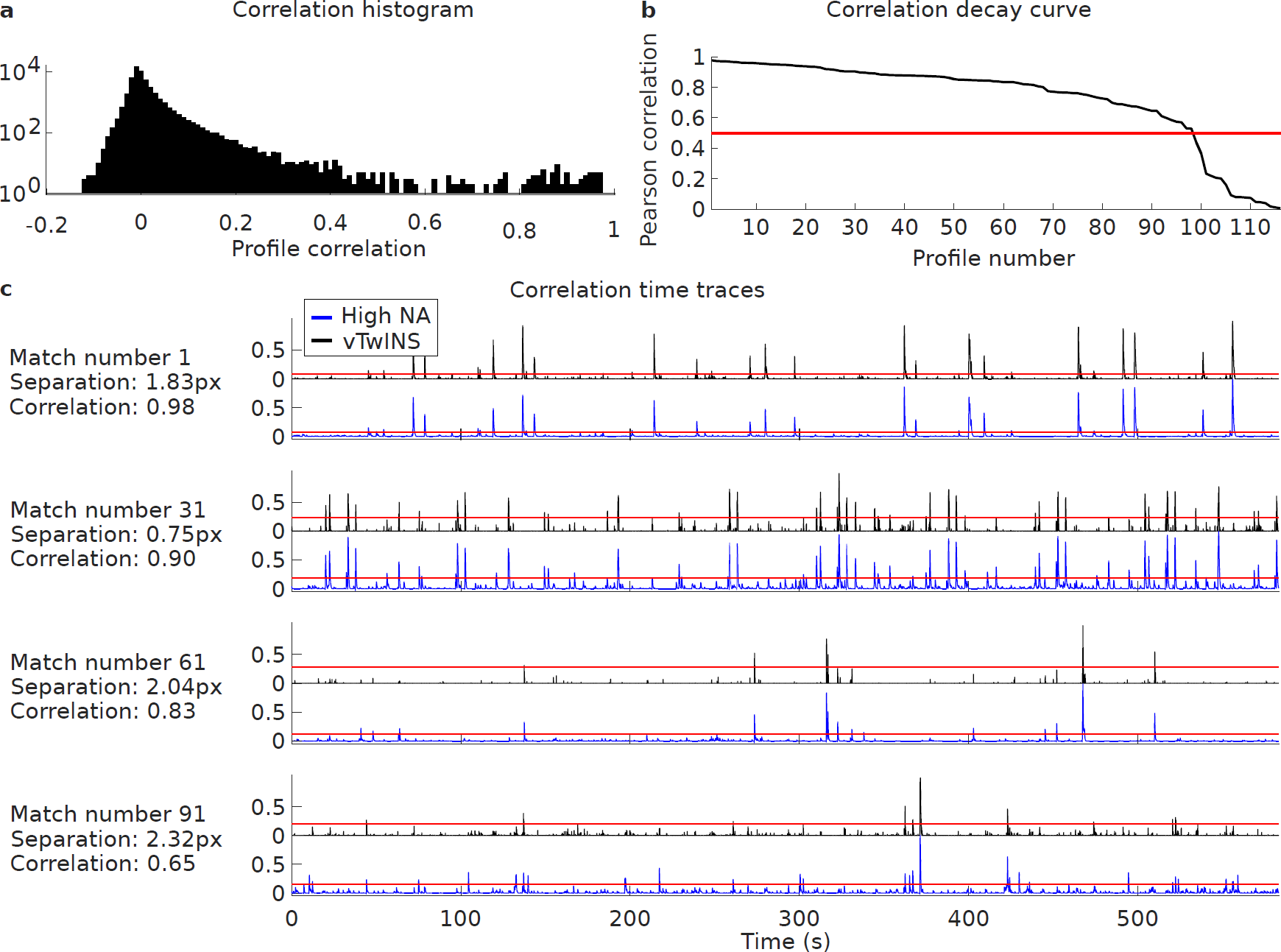
Extended comparison between vTwINS and high-NA spatial profiles and time traces. (a) Histogram of correlations between vTwINS and high-NA time traces. The majority of correlations cluster tightly around zero, and the tail of outliers indicates the correlations for paired traces. (b) Pearson correlations remain high for the top 98 paired profile traces before falling sharply when no more good pairings remain. The red line indicates the ρ = 0.5 cutoff for determining a pairing. (c) Four examples of paired traces for both vTwINS imaging (black lines, top of each pair of plots) and high-NA imaging (blue lines, bottom of each pair of plots). Red horizontal lines indicate the 3σ threshold for significant transients.

### Large Scale Recording in Mouse Hippocampus

As a more challenging application of vTwINS, we recorded and demixed activity from the CA1 region of mouse hippocampus. In this region, neuronal cell soma are densely packed in a well-defined layer; this will produce high spatial overlap in vTwINS data. To induce activity in CA1, water-restricted mice were trained to run down a linear track in a virtual reality system [23] for water rewards (see Methods). Images were collected over a 14 minute session in a 470 *μ*m × 470 *μ*m area with a 35 *μ*m long vTwINS PSF (FWHM, 45 *μ*m 1/e full-width, non-inverted V) at 30 Hz (Supplementary Fig. 8, Supplementary Video 5-6). CA1 recordings were processed and analyzed using the same pre-processing and SCISM demixing as described for the V1 data (Fig. 6a).

**Supplementary Figure 8:**
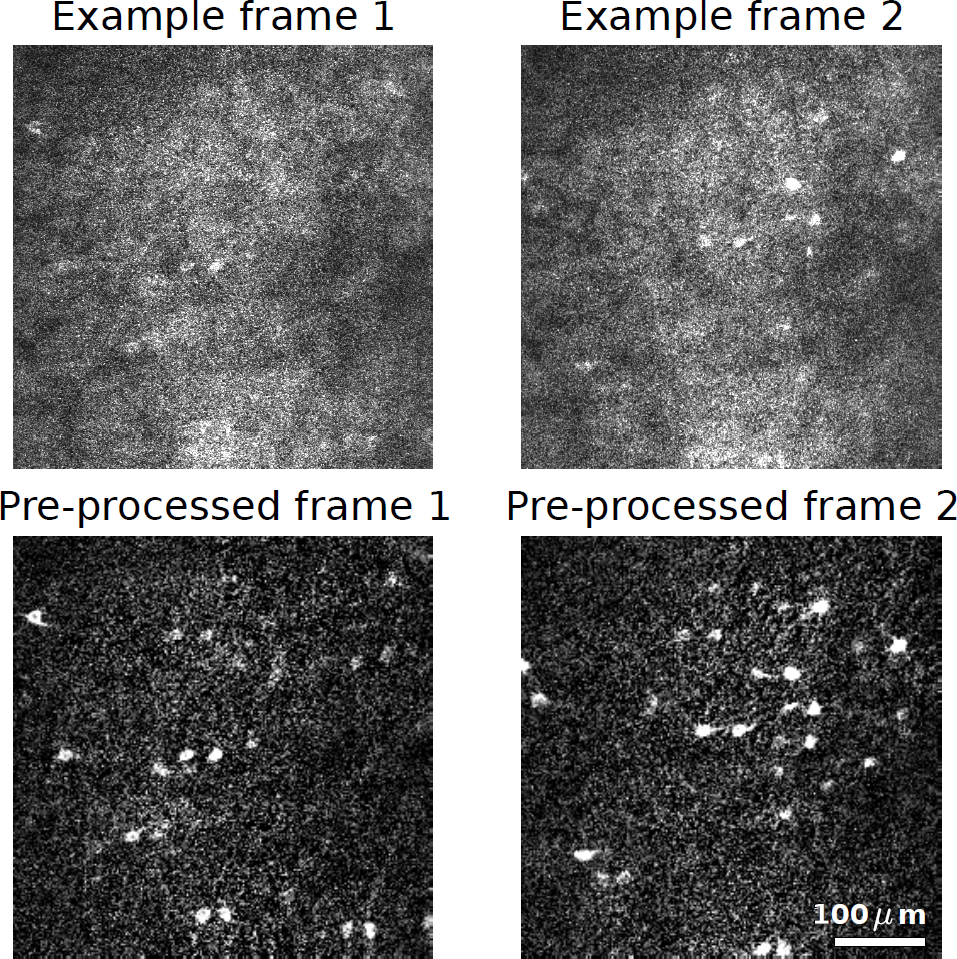
Example frames of full FOV vTwINS data acquired from CA1. Top: two examples of vTwINS images from V1. Bottom: Corresponding pre-processed images (5-frame temporal average and two-fold spatial binning) with background-subtraction. Pre-processing makes active pairs of neuronal images more apparent.

The calculated the 3D positions for each of the 882 spatial profiles found using SCISM span the entire axial range of the PSF (Fig. 6a,b). Interestingly, the tendency for shallower neurons towards the center of the FOV and deeper neurons towards the edges of the FOV, indicates that the vTwINS spatial profiles are capturing the curvature of CA1 (Fig. 6a). The time traces for each of the spatial profiles (Fig. 6c, Supplementary Fig. 9, Supplementary Fig. 10) demonstrate how SCISM selects more active spatial profiles first. We illustrate the range of depths of found spatial profiles as well as example time traces by displaying the spatial profiles situated within the 92 *μ*m × 92 *μ*m white box of Figure 6a and their time traces (Fig. 6b,d). The inferred 3D location of the spatial profiles (Fig. 6b,d) were compared to the anatomical z-stack; the calculated positions correspond well to neurons visible in the anatomical images (Supplementary Fig. 11).

**Figure 6:**
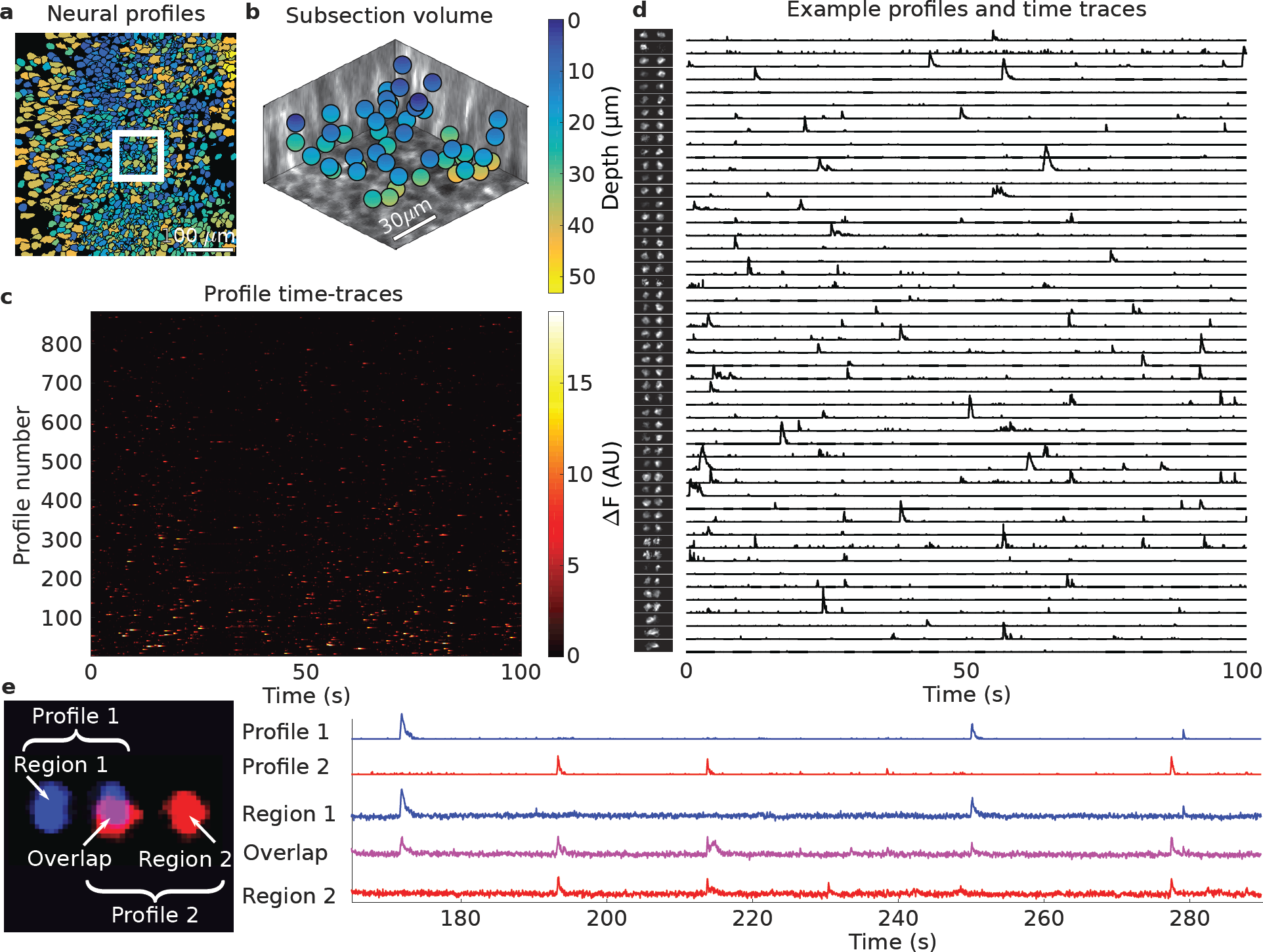
Demixed spatial profiles and calcium activity in mouse hippocampus. (a) Full set of spatial profiles, color-coded by depth, show more overlap in CA1 than in cortical recordings. (b) 3D locations of the spatial profiles from the white box in (a) show found spatial profiles at various depths. (c) Time-traces of spatial profiles in (a) show sporadic activity in the 0-100s time interval.(d) Example subset of spatial profiles (chosen from the white inset box in (a) and sorted by depth) and corresponding time traces show rich activity patterns. Note that some profiles have very sparse activity and do not contain transients in the displayed 100 s range. (e) Example demixed spatially overlapping profiles. Profile 1 (blue) and Profile 2 (red) spatially overlap yet have demixed time traces (right). Averaged raw fluorescence traces from pixels in the overlapping region (Overlap) are a linear combination of the traces from Profile 1 (Region 1) and Profile 2 (Region 2). The Region 2 time-trace also contains a transient from yet another profile at 230 s.

**Supplementary Figure 9:**
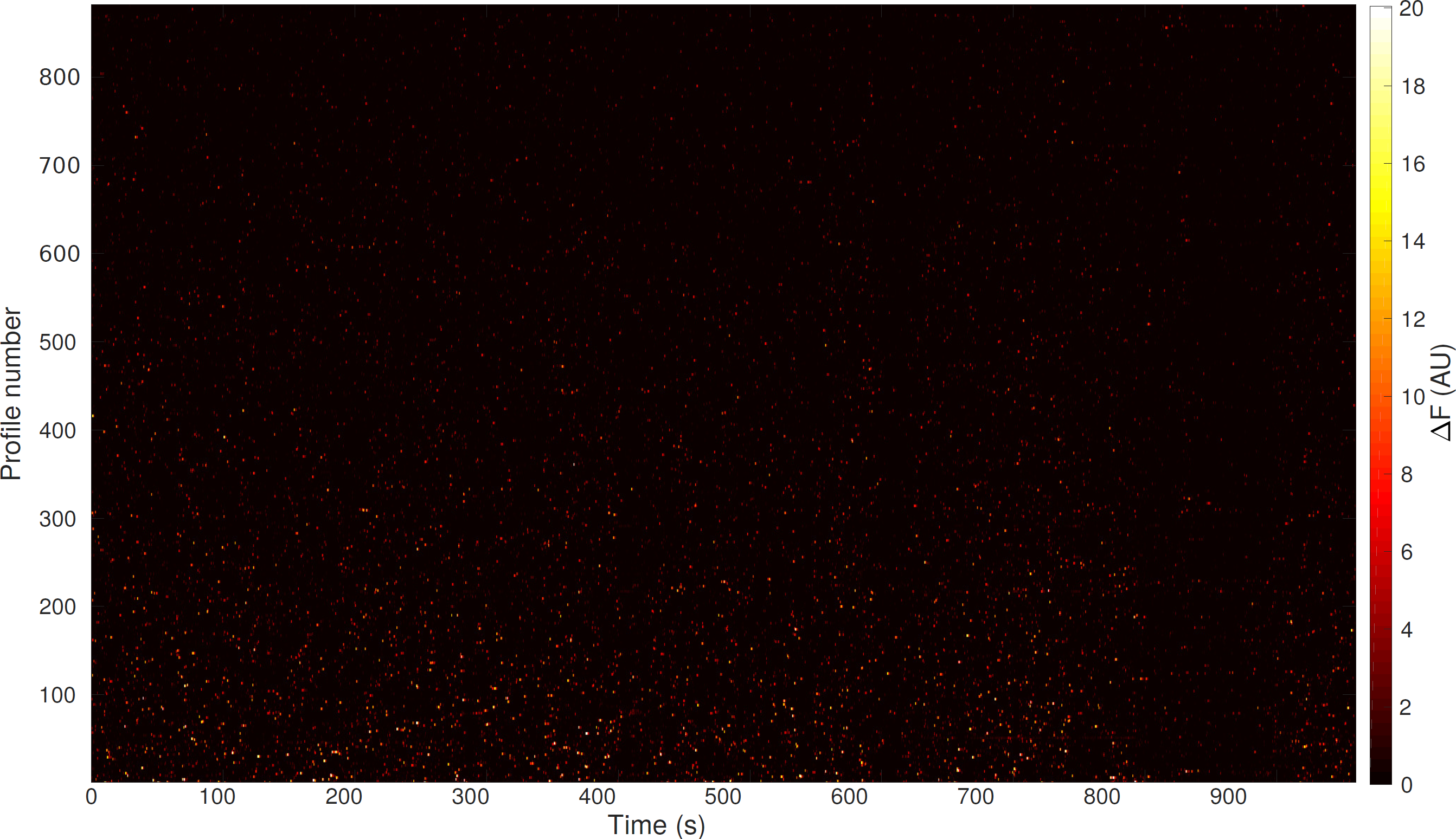
1000 second time traces for all demixed CA1 spatial profiles.

**Supplementary Figure 10:**
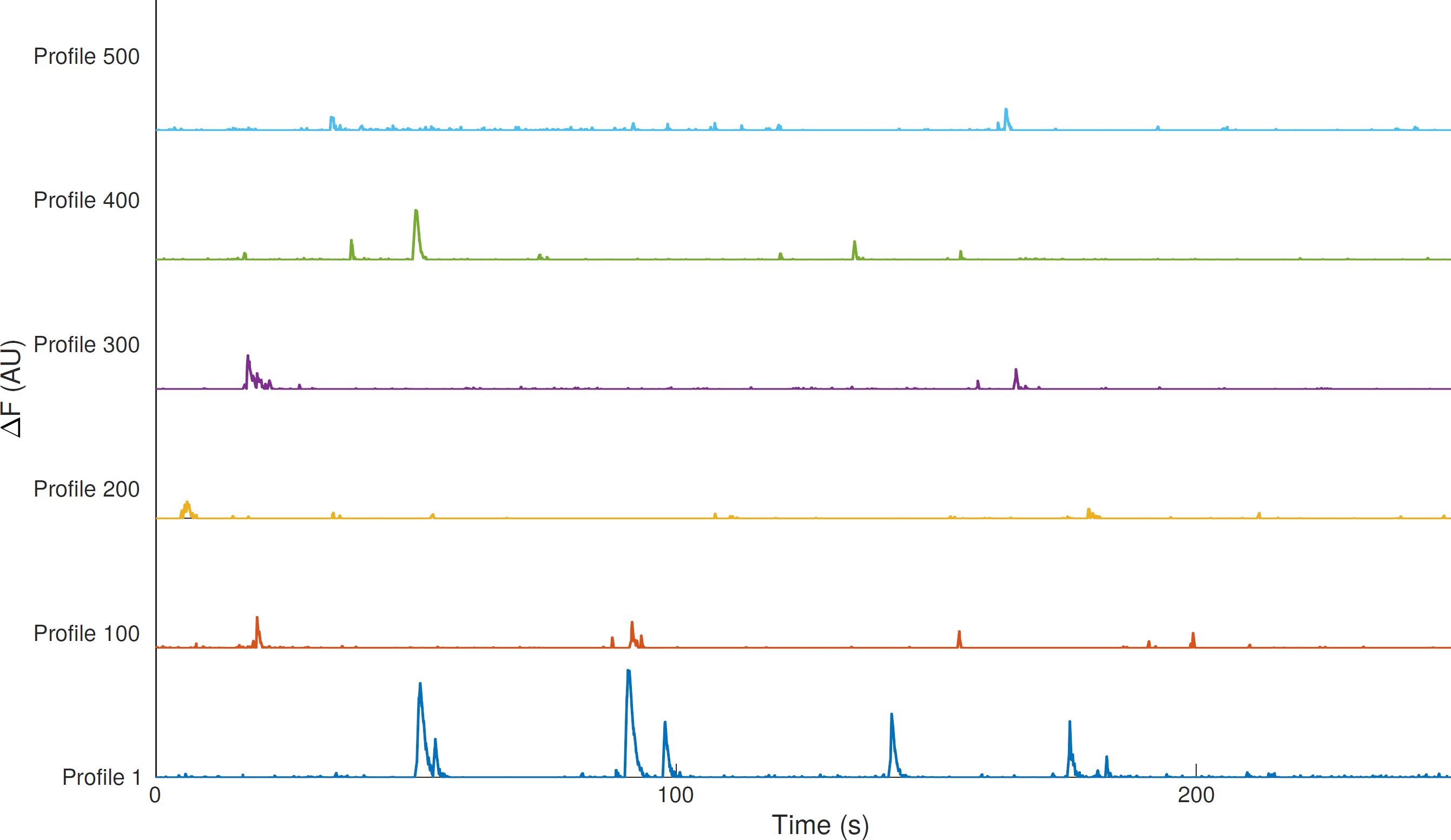
Example time traces for select demixed CA1 spatial profiles.

**Supplementary Figure 11:**
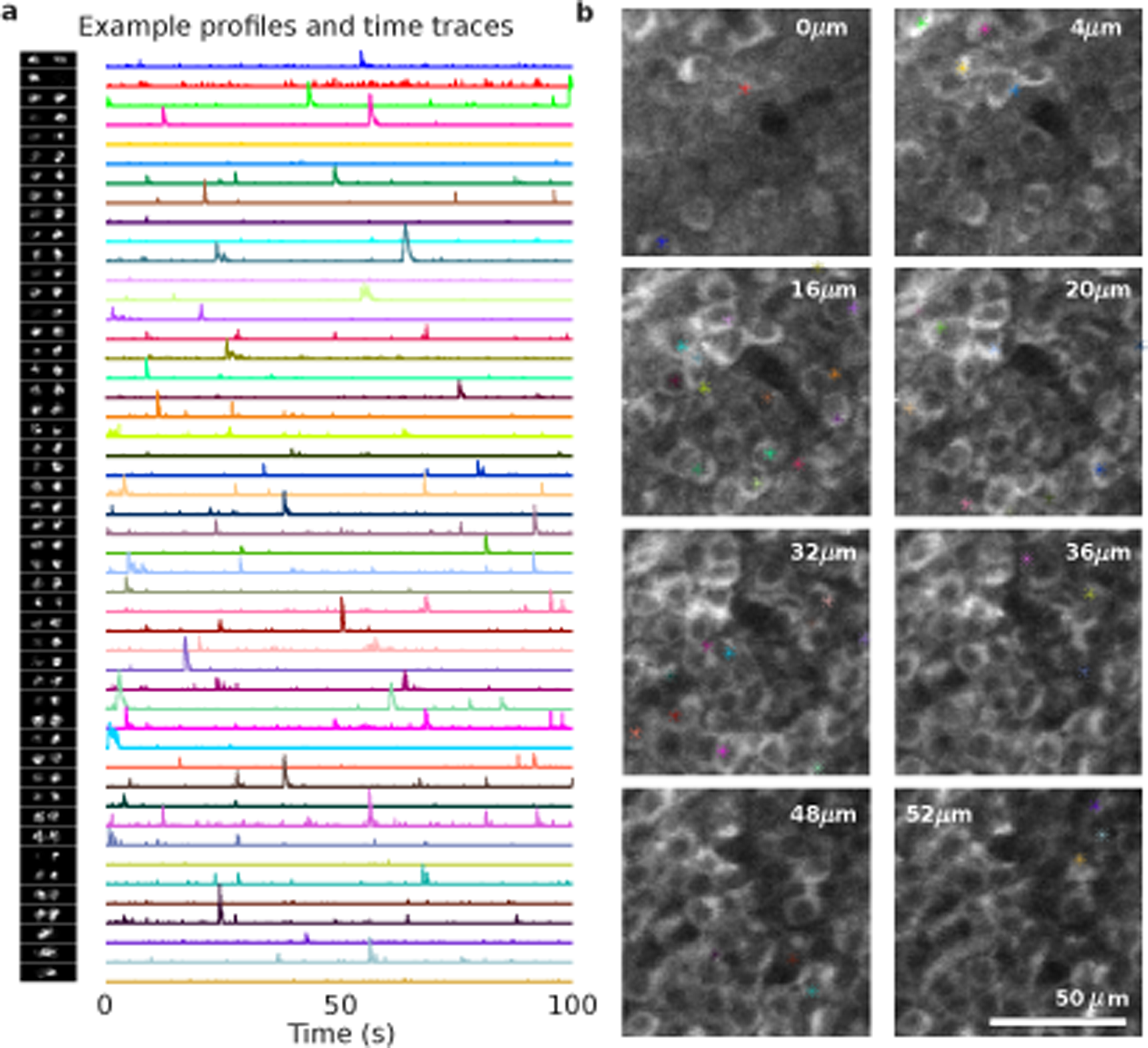
Anatomical z-stack comparison of found spatial profiles in CA1. (a) Example spatial profiles and time traces from CA1. (b) Corresponding location of activity in the anatomical z-stack. White numbers at the top of each image indicates the relative depth in the z-stack.

Despite the highly overlapping spatial profiles due to both the vTwINS PSF and the high overall neural density, vTwINS SCISM successfully demixed spatial profiles in CA1. Fluorescence time courses in different regions of two overlapping spatial profiles illustrate the demixed time traces (Fig. 6e). The trace from the overlapped region of the two cells contains transients from both non-overlap regions, while the demixed traces contain only the single-profile region activity. Interestingly, one transient at 230 s in Region 2's trace is missing from the overlap trace, indicating that this transient originated from a third profile and was successfully demixed in profile 2's time trace.

## Discussion

We have demonstrated that vTwINS can successfully record neural activity from 50 *μ*m thick volumes of the awake mouse brain without a reduction in imaging frame-rate. The novelty of vTwINS primarily lies in using a “V”-shaped stereoscopic PSF to both ensure distinct spatial profiles and encode depth information, and in using priors on the expected spatial profiles and the sparsity of neural activity to motivate a greedy demixing algorithm.

Early strategies for large scale recording using calcium imaging were largely based on the idea of using the spatial resolution of the optical instrumentation to ensure that the fluorescence from individual neurons was collected into a set of voxels that form largely independent, disjoint, sets for each cell. Spatial separation was the basis for hand selection of neural regions of interest (ROIs), which have been widely used as a mask for extracting the time traces of individual active cells. In practice, however, perfect separation of adjacent cell signals into disjoint sets of voxels has been difficult to achieve when expression of the calcium indicator is dense. As a result, demixing algorithms [16, 17, 22] have been developed to identify the spatial profiles corresponding to different cells; these algorithms assume the signal in individual voxels might have contributions from more than one active neuron. vTwINS (and also a recent multi-plane technique [24]) take this mixing assumption as a starting point for the development of the optical instrumentation. The use of a V-shaped PSF in vTwINS increases signal mixing in individual voxels, but it also ensures that each neuron will have a unique spatial profile that can be efficiently used in a co-designed demixing algorithm to extract the time traces of individual cells (and also their location in the volume). We anticipate that this strategy in which optical instrumentation and demixing algorithm are co-designed for large scale recording may generalize to other excitation geometries (e.g. 3 beams, multiple objectives).

vTwINS required the ability to seek specific spatial profile shapes in the recorded movies while maintaining flexibility to reasonably adjust to the particulars of any given dataset. SCISM permitted the specification of these shapes as guides to locate relevant activity while still balancing the general expected temporal statistics of neural activity. Current automated methods do not use such detailed spatial information, focusing instead on temporal demixing [25-29] while imposing no spatial constraints [16, 17] or utilizing generic locality assumptions (i.e. spatial profiles must be fully contained in a constrained region) [22, 30]. The approach in [30] is most similar to ours and incorporates pursuit-type methods within a dictionary learning framework, but does not leverage detailed shape information the way SCISM does. The ability of SCISM to adapt profiles to the data also differentiates it from standard matching pursuit-style algorithms [31-33], which assume a fixed dictionary of features. Although we designed SCISM to seek features specific to vTwINS imaging, it can easily accommodate other spatial profile shapes so as to be applied to non-vTwINS imaging methods.

Additional work can further optimize vTwINS for other applications. In particular, it would be useful to explore how the vTwINS PSF parameters (length, angle between arms, separation distance, beam type) impact imaging in different conditions. Background fluorescence increases with the length of the vTwINS PSF, limiting axial elongation. For neocortical and hippocampal imaging of GCaMP6f under the Emx1-Cre driver, which provides dense labeling of excitatory neurons, we found good performance with ≈10x PSF elongation (5 *μ*m versus 50 *μ*m) despite high neuropil background. In more sparsely labeled tissues, like those provided by Cre driver lines for inhibitory neurons or excitatory neuron subtypes, longer axial extensions are possible. We anticipate that vTwINS might work particularly well with a nuclear localized GCaMP [34], which would significantly reduce background fluorescence from neuropil and improve SNR. The angle between arms and the separation distance influence the available imaging FOV and the required objective lens NA. For brain regions with limited optical access (e.g. hippocampus [23] or MEC [35]), smaller angles between arms or separation distance may be necessary. The choice between Bessel beams and Gaussian beams requires additional study. Bessel beams offer flexibility in controlling the axial profile and lateral resolution [36]. Gaussian beams, while less flexible, are simpler to implement and have higher two-photon excitation. Finally, in applications where lower imaging framerates are acceptable, it should be possible to combine vTwINS with sequential plane imaging (e.g. remote focusing [11], or liquid lens [6]), and image several thin volumes sequentially to image a thick volume.

Motion correction in vTwINS can potentially be compromised due to the elongated PSF reducing high spatial frequencies. In our recordings, however, sufficient high spatial frequencies (vasculature in visual cortex and *stratum oriens* in CA1) facilitated accurate corrections of motion artifacts. For brain regions without distinct high frequency features, expression of a nuclear localized probe tagged with a red fluorophore may be used for accurate motion correction. One unexplored potential use of vTwINS is axial motion correction, which is impossible in single plane TPM. In single plane TPM, axial drift can cause loss of identified neurons over long periods of imaging from the FOV. In vTwINS, each axial position has a unique background shape, potentially allowing the axial position of the imaging volume to be tracked over time. Additionally, vTwINS' axial extension ensures that the vast majority of imaged neurons remain within the imaging volume despite axial drift.

One concern of all large scale TPM calcium imaging methods is photodamage due to the excitation laser, both linear and nonlinear. Nonlinear photodamage in vTwINS as compared to standard TPM is reduced due to both beamsplitting [37] and a lower peak intensity of the axially extended PSF (10x reduction in peak intensity from 50 *μ*m long Gaussian vTwINS beam to a 5 *μ*m long high-NA Gaussian beam). Even with the higher laser power used in vTwINS, the peak intensity is lower than those used in conventional TPM due to the much larger excitation volume. Photodamage due to brain heating, however, should be limited to 200 mW average power at 920 nm [38]. In our setup for vTwINS, we were primarily limited by tissue heating and limited average power to 100 mW per excitation beam.

## Methods

### Microscope Design

The vTwINS microscope was modeled in ZEMAX (Zemax LLC) and custom MATLAB (Mathworks) scripts. The microscope (Fig. 1c) was constructed as a modification of a resonant scanning two-photon microscope. A beam shaping module to produce the V-shaped PSF for vTwINS was designed to be inserted between the laser and microscope. This strategy was used so that the module could, in principle, straightforwardly be adapted for any existing standard two-photon microscope. The beam-shaping module consisted of three optical paths that could be switched via flip-mount mirrors between: 1) a standard high-NA path for standard two-photon imaging, 2) a vTwINS path using low-NA Gaussian beams, or 3) a vTwINS path using Bessel beams.

The collimated Gaussian laser beam entering the beam-shaping module had a measured knife-edge width (10/90 percent) of 1.3mm which corresponds to a 1/*e*^2^ diameter of 2 mm. The high-NA path consisted of a 2.5x beam expander (AC254-40-B and AC254-100-B, Thorlabs). The Gaussian vTwINS path consisted of a 0.3-1.2x variable telescope (G06-203-525 AC 140/31,5 Linos, LC1120 and AC254-125-B, Thorlabs). The Bessel vTwINS path consisted of an axicon and achromat lens pair (179.2° BK7 Axicon, Altechna and AC254-200-B, Thorlabs) to generate the ring-shaped excitation for the Bessel beam. The specific choice of axicon and achromat lens pair was based on tradeoff between lateral resolution and two-photon excitation efficiency. For the Bessel beams to be correctly formed within the sample, the rear pupil of the objective needs to be illuminated with well focused annuli of light. For this reason, the back aperture of the objective is conjugate to the achromatic lens front focal plane of the Axicon-Achromat pair. If collimated, parallel beams are used, the two branches of the PSF form a X-shape. The PSF V-shape was obtained by introducing a slight beam convergence at the objective back-aperture created and tuned by a 1x telescope (2x AC254-100-B, Thorlabs). When the vTwINS modalities were used, the beam was split in two parallel beams with a half-wave plate and a Calcite beam displacer (AHWP05M-980 and BD27, Thorlabs). The half-wave plate was oriented such that the fluorescence intensities of the two images are equal. The birefringent beam displacer was mounted in a rotation mount and oriented such that the two beams lie in a plane perpendicular to the resonant (fast) scanning mirror axis of rotation. This is to guarantee that the two images formed of a fluorescent object lie on the same scanned line. A pair of BK7 windows mounted on orthogonal rotation axes was used to adjust and center the lateral position of the beams on the scanners. The beams separation (2.7 mm out of the Calcite beam displacer) was further reduced using a 0.8x telescope (AC254-100-B and AC254-80-B, Thorlabs). This specific choice, in combination with the magnification of the microscope (X3.75) and the 12.5 mm focal length of the water immersion Nikon objective resulted in an angle of 43° between the two branches of the PSF. This choice of angle resulted in an accurate axial localization of the cell bodies (Supplementary Fig. 1). When the high-NA path was used for conventional two-photon imaging, the half-wave plate was rotated to zero the power of one of the emerging beams, and the two BK7 windows were oriented to center the remaining beam on the optical axis of the microscope.

A Ti:Sapphire laser (Chameleon Vision II, Coherent) at 920 nm was used for two-photon excitation, and dispersion compensation in the laser was adjusted to maximize the two-photon signal. A Pockels cell (Model 350-80 with 302RM driver, Conoptics) was used to modulate laser intensity and a half-wave plate plus polarizing beamsplitter cube (Thorlabs) was used to adjust the maximum laser intensity. The two-photon microscope body consisted of a resonant scanning head (6215/CRS 8 kHz, Cambridge Technologies), a 100 mm *f*-θ scan lens (4401-464-000, Linos) and a 375 mm achromat pair tube lens (2x PAC097, Newport), and an objective lens (N16XLWD-PF, Nikon [39]). The excitation and emission were separated by a shortpass dichroic (T680-DCSPXR-UF3 52 mm × 75 mm × 3 mm, Chroma), and the collection optics (ACL7560-A, LC1611-A, ACL25416UA, Thorlabs) focused the emitted light onto two PMTs (H10770PA-40, Hamamatsu), separated into red and green channels (FF555-Di03-40x54, FF01-720/SP-50, FF02-525/40-32, FF01-593/40-32, Semrock). The PMT signal was amplified with an 80MHz preamplifier (DHPCA-100, Femto) and digitized with a FPGA (NI PXIe-7961R and NI 5732 DAQ, National Instruments). Scanning and data acquisition were controlled with Scanimage 2015 (Vidrio). Average power during vTwINS data acquisition varied between 150 mW and 200 mW at 920 nm, and average power during high-NA acquisition was between 50 mW and 70 mW at 920 nm. Images here were typically acquired at 30Hz with an image size 512x512 pixels at with a 90% spatial cutoff, corresponding to an image size of 470 *μ*m x 470 *μ*m (2.8 zoom) or 550 *μ*m x 550 *μ*mm (2.4 zoom). Nearly simultaneous calcium imaging using rapid switching between vTwINS excitation and the traditional focused high-NA Gaussian PSF was performed using an alternate optical setup (Supplementary Fig. 6). A galvanometer (6210H, Cambridge Technologies) was used to select between high-NA and vTwINS paths, which were recombined downstream with a (50 *μ* 88° optical) offset. A modified Scanimage analog control was used to switch between the two paths at every frame. For each modality, images were acquired at 30 Hz with a 512x256 pixel image size.

### Transgenic Mice

All experimental procedures were approved by the Princeton University Institutional Animal Care and Use Committee. Transgenic GCaMP6f-expressing mice were produced by crossing Emx1-Cre (*B*6.129 *S*2–*Emx*1^*tm1(cre)Krj*^/*J*, Jax #005628), CaMK2-tTA (B6.Cg-Tg(Camk2a-tTA)1Mmay/DboJ, Jax #007004) and TITL-GCaMP6f (Ai93; *B*6.Cg-Igs7^*tm93.1(tetOGCaM P6f)Hze*^/*J*, Jax #024103) strains [40]. Male or female transgenics heterozygous for all three genes were used for all experiments.

### Imaging Mouse Visual Cortex

For imaging in mouse visual cortex, mice underwent surgery under isoflurane anesthesia for implantation of imaging windows and head-plates. A 5 mm diameter craniotomy was made over one hemisphere of parietal cortex (centered 2 mm caudal, 1.7 mm lateral to bregma). A custom titanium head-plate and optical window (#1 thickness, 5 mm diameter glass coverslip, Warner Instruments) bonded to a steel ring (0.5 mm thickness, 5 mm diameter, SS316 ring, Ziggy's Tubes and Wires, Inc.) were attached to the mouse's skull with dental cement (Metabond, Parkell). The location of V1 was estimated using a separate wide field imaging microscope to record retinotopic responses in fluorescence activity as the mouse viewed horizontally and vertically drifting bars on a 32” monitor [41]. Boundaries between the primary and secondary visual areas were defined using an automated algorithm to locate reversals in the retinotopic gradients [42]. Five days after surgery, mice were trained to run on a spherical treadmill (8 inch diameter Styrofoam ball) surrounded by a 270° toroidal screen [43]. Visual stimuli were generated using the Psychophysics Toolbox [44-46] and displayed on the toroidal screen using a DLP projection system (Mitsubishi HC3000), consisting of ≈100 randomly placed/oriented Gabor patches, with visual field size 5-10°, updated at 4 Hz. To prevent light from the projected display from entering the fluorescence collection system, the region between the base of the objective lens and the head-plate was light-proofed using a black rubber tube prior to imaging. The rubber tube was glued to a silicone firing and the firing itself attached to the titanium headplate with silicone elastomer (Body Double, Smooth On Inc.). Examples of images from cortical imaging are depicted in Supplementary Figure 2, Supplementary Figure 12 and Supplementary Video (1-4).

**Supplementary Figure 12:**
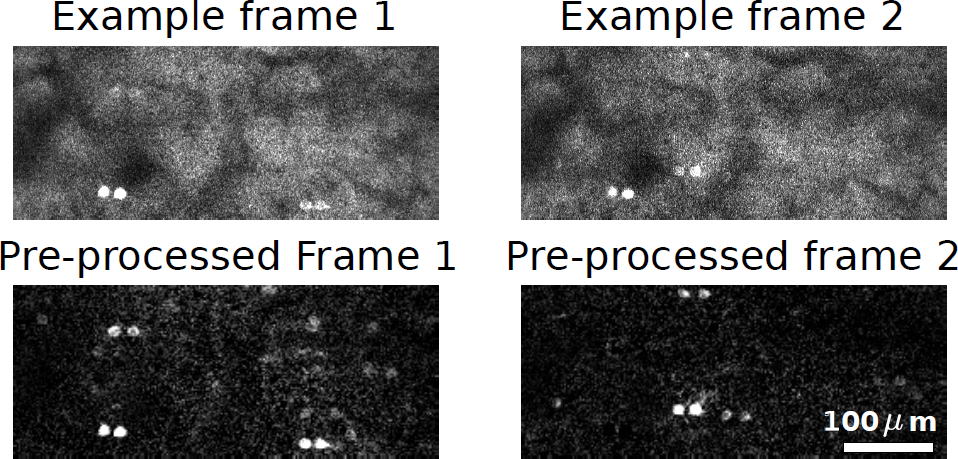
Example frames of half FOV vTwINS data acquired from V1. Top: two examples of vTwINS images from V1. Bottom: Corresponding pre-processed images (5-frame temporal average and two-fold spatial binning) with background-subtraction. Pre-processing makes active pairs of neuronal images more apparent.

### Imaging Mouse Hippocampus

For imaging in mouse hippocampus, mice underwent surgery under isoflurane anesthesia for implantation of an imaging window and a head-plate for head-restraint in virtual reality [47]. Optical access to the hippocampus was obtained as described previously [23]. Briefly, a ≈ 3 mm diameter circular craniotomy over the left hemisphere was performed, centered 1.8 mm lateral to the midline, and 2.0 mm posterior to bregma. The cortical tissue overlying the hippocampus was aspirated, and a circular metal cannula with a #1 coverslip bonded to the bottom was implanted, with a thin layer of Kwik-sil (WPI) between the hippocampus and coverslip. During the surgery, a titanium headplate was attached to the skull with Metabond. After recovery, mice were water restricted for five days and then trained to run on a 4 m virtual linear track using a virtual reality setup similar to that described in Domnisoru et al. 2013 [48]. Visually distinct towers were placed every 1m and 4 *μ*L water rewards given at 1.6 m and 3.6 m down the track. Mice ran on a 6 inch diameter Styrofoam cylinder (The Baker's Kitchen) whose position was detected by an angular encoder. Mice were trained for a 60 minute session per day and were given 1-1.5 mL of water a day total (including behavioral training and supplemental water). The virtual reality projection system was as described previously [43, 47] and controlled with ViRMEn [49]. Lightproofing around the objective was performed as described for experiments in visual cortex. Examples of images from hippocampal imaging are depicted in Supplementary Figure 8 and Supplementary Video (5,6).

### Fluorescent Bead Sample and Measured PSFs

As an initial test of vTwINS we imaged 1 *μ*m green fluorescent beads (L1030, Sigma) embedded in a 1% agarose gel. The beads were embedded at random locations, creating an off-grid set of positions. The exact bead positions were determined via a diffraction-limited two-photon multi-plane volumetric scan (z-stack). vTwINS was then used to image the same volume with a single scan (one image). As shown in Figure 1 each bead appears in the vTwINS projection image as a pair of dots; lines drawn between all pairs are parallel and aligned with the direction of the vTwINS PSF in the sample. The distance between dot pairs varies with the bead's depth in the volume. Using the single vTwINS image, SCISM was used to automatically locate the spatial profiles for each bead. The found profiles were used in turn to infer each bead's 3D coordinates in the sample, using the vTwINS relationship between depth and inter-image distance. Beads at the edge of the imaged volume with only one projection into the vTwINS image were discarded as the depth location could not be ascertained.

Z-stacks of these fluorescent beads (1 *μ*m step size) were taken to measure the vTwINS PSFs for each set of experiments. The axial length of PSFs were measured by averaging the fluorescence intensity at each slice of the z-stack. The averaged fluorescence signal was used to calculate the FWHM and 1/e full-width axial lengths. The PSFs were not measured *in vivo*, although this can be done to correct for index mismatch and scattering if higher accuracy is desired.

### Motion Correction and Pre-processing

All video sequences were first subject to a normalized cross-correlation-based motion correction algorithm. This algorithm, implemented via the template matching function of OpenCV [50], found the best horizontal and vertical shifts for each frame to match a fixed template. The template used was set to the median across frames. Shifts were set to have a maximum allowable value (set to 10 pixels for the V1 data and 15 pixels for the CA1 data). Videos were cropped to remove edge rows and columns with missing data due to shifting.

After motion correction, all data was subject to spatio-temporal averaging as a pre-processing step aimed to improve SNR and run-time of the demixing algorithm. Specifically, five-frame temporal running averages and a two-fold spatial binning were applied to the data prior to running a demixing algorithm. We ran our automated demixing algorithm on the pre-processed vTwINS movies. Although the two-fold spatial binning is not required for demixing, it greatly improved run-time.

### vTwINS Orthogonal Matching Pursuit

In this section, we describe the mathematical details of the vTwINS Sparse Convolutional Iterative Shape Matching (SCISM) demixing algorithm. Let ***Y*** ∈ ℝ*^N×T^* denote the calcium video sequence, ***X*** ∈ ℝ*^N×K^* denote the neural spatial components (spatial profiles), and ***S***∈ ℝ*^T×K^* denote the neural temporal activity traces, where *N* is the number of pixels in each image, *T* is the number of images (or time points), and *K* is the number of neurons. Thus, the columns of ***Y*** represent single frames of the video, the columns of ***X*** represent individual spatial profiles, and the columns of ***S*** represent temporal activity traces of single neurons. We model background activity with a set of *B* background components ***X****_bg_*∈ ℝ*^N×B^* and denote the (inferred) background temporal activity ***S****_bg_* ℝ*^T×B^*.

Our algorithm is designed to exploit *a priori* knowledge of both the spatial profile shapes as well as neural firing statistics. Specifically, the algorithm seeks to factor the full movie matrix ***Y*** into the set of spatial profiles ***X*** and time-traces ***S***such that

1. The sum of outer products of spatial profiles and time traces explains the observed data (***Y*** ≈ ***XS****^T^*).
2. The time-traces ***S*** are sparse in time.
3. The spatial profiles are shaped like pairs of neuronal somata (disks or annuli), offset horizontally by a small separation distance. The dark center in each soma is due to the lack of GCaMP6f in the nucleus.
4. Few latent sources (active neurons) relative to the size of the data generate activity in the observed data, making the fluorescence movie low-rank. This constraint captures the physical density constraints on neuron tissue.

The optimization program that includes all these terms is

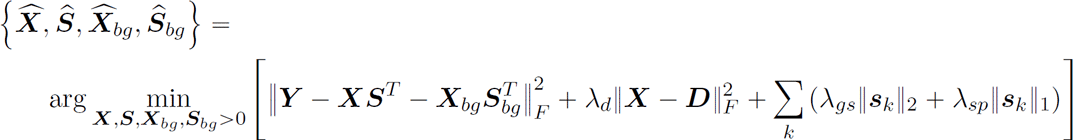

where *s_k_* is the *k^th^* column of ***S***, representing the activity of neuron *k*, 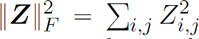 is the squared-Frobenius norm, ***D*** is a matrix whose columns represent all possible expected neural spatial profile shapes, *λ_d_* is the trade-off parameter for penalizing the deviation of spatial profile shapes ***X*** form the idealized shapes in ***D***, *λ_gs_* is the group-sparse penalization parameter for ensuring that not all spatial profiles are active and *_sp_* is the penalization parameter than ensures the time traces are sparse. Each column ***d****_k_* of ***D*** represents the expected spatial profile for a neuron at one volumetric neural location. We set the spatial profiles ***d****_k_* as annuli separated by a depth-dependent distance (Fig. 3a), where the annuli were modeled as the difference of two Gaussian functions, separated by a distance

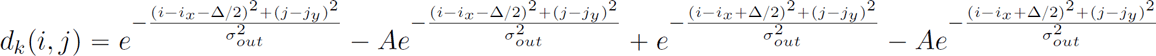

For all datasets analyzed here, the annuli were set to have *σ_out_* = 2 pixels and *σ_in_* = 0.84 pixels, the center amplitude depression was set to *A* = 0.7, and (*i_x_*, *j_y_* simply indicate the pixel which ***d**_k_* is centered around. We used 10 different inter-image distances, Δ, equally spaced between 21.4µm to 92.4µm for full FOV V1 data spanned, 18.4 µm and 51.4 µm for half FOV V1 data, and 14.6 µm to 56.8 µm for full FOV CA1 data. In total, the number of columns of ***D*** is the number of pixels *N* (all potential spatial locations) times the number of inter-image distances *K* (***D***∈R*^N×NK^*). This matrix, however, never needs to be constructed, as any the spatial invariance of the neural profiles permits the use of convolution operations. The parameters used for our analysis reflects the particulars of our microscope setup (i.e. zoom, beam angle setting etc.) and should be modified to fit the expected statistics of any new dataset.

The optimization program in Equation (1) results from modeling the measurement noise as Gaussian, and placing appropriate sparsity- and shape-penalizing priors on the spatial profiles and transients. The measurement model and Gaussian prior over the spatial profiles ***X*** are given by

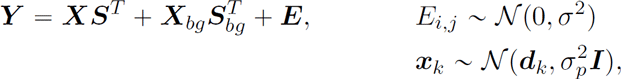

where the non-zero mean of the Gaussian prior over spatial profiles induces the expected spatial structure. The prior over time traces*s_k_*

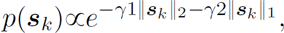

includes two terms penalizing both overall sparsity and group sparsity (each neural trace being a group). In terms of the model parameters, the trade-off parameters in Equation 1 are 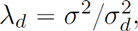 λ_gs_ = γ1σ^2^, λ_sp_ = γ2σ^2^. No specific prior was placed on either the background shape or its temporal fluctuations.

Direct optimization of Equation (1) can be inefficient due to the problem size and the large search space (potential spatial profiles). We thus approximated a solution to Equation (1) with a greedy, iterative approach wherein spatial profiles are sequentially determined. Our method iterates between finding the best element of ***D*** that approximates ***Y*** given the sparsity constraints and updating that profile to the data. The first step sets ***X***=***D*** and solves for the best single trace to approximate ***Y*** (solving the first and third terms). The shape refinement step then uses the first two terms with the newly found time-trace to allow the spatial profile ***x**_k_* to deviate from its mean ***d**_k_*. SCISM is in essence a modification of the orthogonal matching pursuit (OMP) method for greedy sparse signal estimation [31, 51]. Our method extends OMP by including an additional temporal sparsity penalty and a shape refinement step that allows for deviations from the stereotyped neuronal shapes (traditional OMP assumes a fixed dictionary of features).

We initialized our algorithm by estimating the background spatial profile, ***X***_bg_ using the normalized temporal median of the pre-processed motion-corrected video sequence and ***S***_bg_ as its least-squares time course

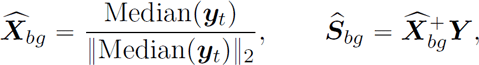

where ***X***^+^ denotes the pseudo-inverse of***X***. In the case of shorter video sequences (20000 frames or less) we only used a single background spatial profile (*B* = 1). For longer video sequences a background spatial profile was added for each 5000 consecutive and non-overlapping frames, which allowed the background to change over the course of the video sequence (e.g. due to slow axial drift). The *residual* movie ***R*** was then initialized at the first step to the median-subtracted full movie ***Y***

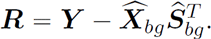

The algorithm (summarized graphically in Fig. 3 and algorithmically in Alg.1) begins each iteration by seeking the stereotyped annuli pair that had the largest correlation with the residual movie ***R***. Specifically, the algorithm seeks the index *k* and the corresponding pair *d_k_* with the largest value *v_k_* calculated as,

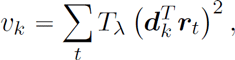

where **T**_λ_ is a soft thresholding function restricted to positive values

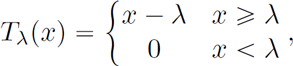

and ***r**_t_* are the columns of ***R*** (the frames of the residual video). *v_k_* estimates the total energy of the estimated time trace *s_k_* that minimized Equation (1) conditioned on *x_k_* = ***d**_k_*, and all past profiles and time traces being fixed. The thresholding operation induces temporal sparsity (the last term in Eqn.(1)), and prevents noise accumulation over long videos from dominating the values of *v_k_*. Thus even very sparsely firing neurons can be identified, provided they fluoresce above the noise floor. Because the noise floor is not spatially constant, we set the sparsity penalization parameter λ to be a function of the local statistics effecting each potential spatial profile shape. Specifically, we set λ to be proportional to the 99*^th^* percentile of the residual projected into the stereotyped shapes,

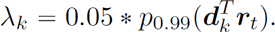

This local parameter setting measures the potential brightness at each location. As brighter locations have higher backgrounds and higher noise levels, λ is thus set higher at these locations.

After calculating *v_k_*, the stereotyped spatial profile 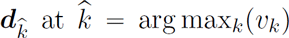 is added to the set of spatial profiles. As ***d**_k_* only approximates the profile shape, a spatial profile that balances the observed data and prior shape information is obtained using a shape refinement step. The shape refinement step estimates ***x**_k_* from ***R*** and ***d**_k_* as

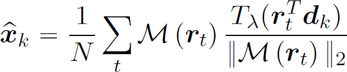

where ***M***(·) is a mask that restricts the averaged frames to the location of ***d**_k_* (thereby preventing spurious activity from across the video from being included in the spatial profile ***x**_k_*). The normalization by the magnitude of ***r**_t_* prevented spurious high activity frames, where the activity may not come from that particular neuron, from dominating the average and corrupting the results. In terms of the original cost function, this essentially prevents contributions from yet-to-be located neurons from in uencing the spatial profile of the current neuron. While SCISM could be modified to refine all past spatial profiles at each iteration in order incorporate the new profile, such an extension is not explored here.

Given the updated spatial profile list, the time traces ***S*** and ***s**_bg_* are obtained via non-negative LASSO

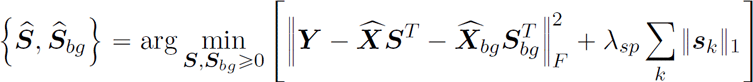

and the residual movie is updated as

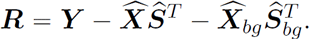

The algorithm then repeats, using the new residual to find the next neural spatial profile, starting again from Equation (2).

We ran SCISM until either a pre-set number of spatial profiles was found, or the activity trace for the most recently found spatial profile was essentially zero. Ideally, however, SCISM would iterate until the recovered spatial profiles no longer resemble neurons. While our neural activity-based criterion attempted to determine if newly found spatial profiles represented neurons, more sophisticated methods would increase accuracy. Since testing if a spatial profile represents a neuron is still an open problem [20], one potential approach is to manually check new spatial profiles as they are found and manually stop when newer profiles are deemed to no longer be capturing neural activity. An example of SCISM processing vTwINS data is provided in Supplementary Video 7.

Once the algorithm is ended, the full-temporal resolution time-traces is obtained via non-negative LASSO (Eqn. (6)) with the non-temporally averaged data in place of ***Y***.

#### Algorithm 1

SCISM algorithm for locating pairs of neuronal images in volumetric calcium data.

~~~
1: Set λ_1_,λ_2_ and K or *s*
2: Set m = 1
3: Initialize 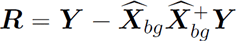
4: 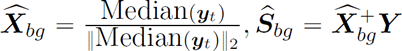
5: **repeat**
6:  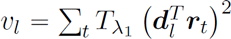
7:   k = arg max_l_ v_l_
8:  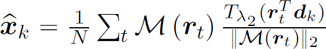
9:  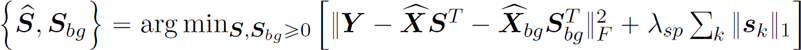
10:  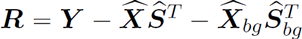
11:  m = m + 1
12: 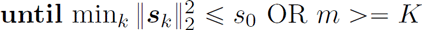
13: 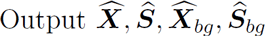
~~~

To improve the computational efficiency of our method, we introduced two optional modifications to the algorithm's order of computations. First, because inner products of distant spatial profile shapes are nearly independent, multiple new spatial profiles can be selected at each iteration by seeking multiple, well-separated, local maxima of *v_t_*. Second, calculating all inner products with the residual at each iteration can be computationally expensive (essentially *K* 3D convolutions between each ***d**_k_* and the data). For small-to-medium sized datasets, we offset some of the computational burden, at the cost of additional memory, by using the linearity of the inner product. Using the reformulation

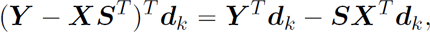

the algorithm could pre-calculate the inner products with the data (***Y ∗ d_k_***) and the spatial profiles (***X^T^d_k_***), and the inner products with the residual were then calculated via a small number of outer products ***S***(**X**^T^*d_k_* and a subtraction operation. The computational savings of this reorganization, however, were diminished for larger datasets where memory allocation became as burdensome as calculating convolutions. SCISM was implemented in MATLAB and made use of the TFOCS library [52] to solve the weighted, non-negative LASSO optimization step. Typical analysis ran at a rate of approximately 20 s per profile found.

### vTwINS and High-NA Spatial profile Registration

High resolution anatomical z-stacks (median of 200-300 frames per slice at 2.5-4 *μ*m slice separation taken with the high-NA beam path) were taken for each vTwINS imaging volume to align the vTwINS spatial profiles to anatomical positions. Alignment between the anatomical z-stack and the vTwINS imaging volume was performed in two steps: 1. The 3D position of cells was estimated to their position within the vTwINS volume. 2. The estimated 3D positions were offset to the anatomical volume. First, the centroids of each half of the spatial profile were used to calculate the 3D cell position via *d* = 0.5(△–△_min_)/tan(θ), where △_min_ is the minimum inter-beam distance of the PSF and θ is the beam angle from the axial direction. A correction to the xy position was made for any differences in θ between the two halves of the vTwINS PSF. Second, a 3D offset between the estimated positions and anatomical z-stack positions was either automatically or manually calculated. For automatic alignment, the anatomical stack was first deconvolved (Lucy-Richardson) with the high-NA PSF and then convolved with the vTwINS PSF. A 3D cross-correlation was then calculated between the convolution stack and the median vTwINS image and the peak of the cross-correlation was used as the offset between the vTwINS images and the anatomical z-stack. For manual alignment, highly active cells with similar cell shapes between the vTwINS spatial profiles and high-NA anatomical z-stacks were located manually and used to estimate the offset between the vTwINS images and anatomical z-stack.

For simultaneous vTwINS and conventional TPM imaging, neural activity was independently extracted from raw images with separate analyses. Neural activity underlying calcium dynamics for conventional TPM was estimated using the Constrained Non-negative Matrix Factorization and deconvolution algorithm (CNMF) to demix contributions from possibly overlapping cells [22, 53]. Spatial profiles extracted using CNMF were manually selected for regions that approximated a cell shape (roughly circular, 10–15 µm in diameter). To compare number of spatial profiles between imaging modalities, spatial profiles from either methods were only included if their center position was within 20 pixels (18 µm) of the x (fast-scanning) edge of the acquisition region. This is to prevent bias from clipping half of a single vTwINS profile near the edges of the image.

Spatial profiles and time traces extracted using vTwINS OMP and CNMF were paired off by their normalized time trace Pearson correlation (Supplementary Fig. 7), subject to the constraint that the extracted spatial profile center positions were within 5 pixels (4.5 µm) in the y (slow-scanning) direction and 40 pixels (37 µm) in the x (fast-scanning) direction. This distance is roughly equal to half the maximum separation distance between vTwINS spatial profile image pairs, which does not restrict pairing of CNMF spatial profiles to vTwINS spatial profiles with a single blocked beam. Spatial profiles and time traces were paired off until the correlation dropped below a 5σ excess of the average correlation between any two time traces. Only high activity cells with >1 statistically significant transient/min [54] were included for this analysis. A transient was considered statistically significant if its peak was >3σ above the average noise levels.

## Acknowledgments

We would like to thank Cristina Domnisoru, Ryan Low and Ben Scott for their insightful thoughts and comments. We also thank Jan Homann for his assistance using Psychophysics Toolbox. This work was supported by NIH grants R01MH083868 and U01NS09054, and the Simons Collaboration on the Global Brain. A.C. was supported by an NIH NRSA Training Grant in Quantitative Neuroscience (T32MH065214). J.W.P was supported by grants from the McKnight Foundation, Simons Collaboration on the Global Brain (SCGB AWD1004351) and NSF CAREER Award (IIS-1150186).

## Author Contributions

D.W.T. conceived the project. A.S. and S.Y.T. designed and constructed the vTwINS microscope. S.A.K. and J.L.G. performed the surgery on the mice. A.S. trained the mice and performed the imaging experiments. A.S.C. and J.W.P. designed the SCISM algorithm. A.S.C. implemented SCISM and applied the method to the vTwINS data. A.S. and A.S.C. performed the analysis on the results. A.S., A.S.C., J.W.P., and D.W.T wrote the manuscript, with comments and contributions from all authors. J.W.P and D.W.T supervised the project.

## Supplementary Figures and Videos

Supplementary Video 1: Video of motion-corrected vTwINS images acquired from V1 and used in Figure 4.

Supplementary Video 2: Video of pre-processed vTwINS images acquired from V1 and used in Figure 4. Images were pre-processed images (5-frame temporal average and two-fold spatial binning) with background-subtraction (median of movie).

Supplementary Video 3: Video of motion-corrected vTwINS images acquired from V1 and used in Figure 5.

Supplementary Video 4: Video of pre-processed vTwINS images acquired from V1 and used in Figure 5. Images were pre-processed images (5-frame temporal average and two-fold spatial binning) with background-subtraction (median of movie).

Supplementary Video 5: Video of motion-corrected vTwINS images acquired from CA1 and used in Figure 6. Images were averaged with 5 frames temporally.

Supplementary Video 6: Video of pre-processed vTwINS images acquired from CA1 and used in Figure 6. Images were pre-processed images (5-frame temporal average and two-fold spatial binning) with background-subtraction (median of movie).

Supplementary Video 7: Video corresponding to SCISM demixing for V1 data Figure 4. The upper left portion of the videos shows the spatial locations of all profiles found in all iterations thus far. Each pair of circles with a connected line show the pair of images comprising the found profiles. The upper right portion of the video shows the pair locations (circles with connected lines from the upper left portion) super-imposed on the mean-to-variance ration of the movie (a simple measure of activity). This demonstrates that pairs of activity locations f similar strength are picked up. The bottom portion of the video shows the time traces for the set of profiles found at the current iteration.

